# Light on Broken Networks: Resting-State fNIRS as a Tool for Connectivity Mapping

**DOI:** 10.64898/2026.03.06.710143

**Authors:** Foivos Kotsogiannis, Michael Lührs, Geert-Jan M. Rutten, Andrew T. Reid, Sabine Deprez, Maarten Lambrecht, Wouter De Baene, Charlotte Sleurs

**Author notes:** Corresponding author: Charlotte Sleurs, Department of Cognitive Neuropsychology, Tilburg University, Tilburg, The Netherlands.

## Abstract

Resting-state functional connectivity (RSFC) and networks (RSNs) provide insight into large-scale brain organization and its disruption in neurological disease. RSNs are most commonly assessed using fMRI, yet its translational use is constrained by high cost, motion sensitivity, and limited feasibility for repeated measurements. Functional near-infrared spectroscopy (fNIRS) offers a portable alternative, but its reliability for RSFC and RSN mapping remains insufficiently established. Near whole-head fNIRS data and fMRI-BOLD signals of corresponding cortical regions were extracted, based on which RSN organization was compared across two independent cohorts of 31 participants each. Cross-modal convergence and divergence were assessed using bivariate and partial correlations across multiple network levels. Edgewise analyses revealed substantial modality differences with bivariate correlations (50–61% of edges), which were markedly reduced using partial correlations (<3%). Group-level connectivity patterns showed moderate cross-modal similarity (*r* ≈ 0.37). At nodal level, net strength, local efficiency, and path-length differed substantially between modalities, while normalized strength and assortativity were largely comparable. Across nodes, group-level graph-metric distributions were broadly similar for normalized strength, assortativity, local efficiency, and path length (*rho* ≈ 0.27–0.5). At network-level, fNIRS-derived modules significantly overlapped with fMRI modules, particularly based on bivariate correlations, identifying default mode, attentional, executive, salience, sensorimotor, and visual networks (Jaccard ≈ 0.27–0.5). Overall, fNIRS captured key features of large-scale RSFC and RSN organization observed with fMRI, supporting meaningful cross-modal correspondence and translational utility. While partial correlations enhanced edge-level agreement, they attenuated nodal and modular recovery, suggesting greater suitability for targeted connectivity analyses rather than whole-network characterization.

## 1. Introduction

The human brain consists of complex, interconnected networks that can be characterized in terms of both structural and functional connectivity (Babaeeghazvini et al., 2021; Cabral et al., 2017; Erol and Hunyadi, 2022; van den Heuvel and Hulshoff Pol, 2010). Structural connectivity refers to anatomical connections, which remain relatively stable across time (Babaeeghazvini et al., 2021; Cabral et al., 2017; Park et al., 2008; Soldozy et al., 2020). Functional connectivity (FC), by contrast, is dynamic, reflecting the temporal coherence in neural activity across anatomically separated regions (Babaeeghazvini et al., 2021; Erol and Hunyadi, 2022; van den Heuvel and Hulshoff Pol, 2010), and can occur between areas without direct anatomical projections (Biswal et al., 2010).

Functional magnetic resonance imaging (fMRI) has played a key role in mapping functional connectivity, particularly through resting-state paradigms where participants are not performing explicit tasks (Smitha et al., 2017; van den Heuvel and Hulshoff Pol, 2010). The relevance of resting-state (rs) fMRI was first demonstrated in the landmark study by Biswal et al. (1995), which revealed coherent resting-state functional connectivity (RSFC) within the motor network. Resting-state paradigms are straightforward to implement (O’Connor and Zeffiro, 2019), require minimal participant engagement and can be applied to individuals unable to perform task-based paradigms (Canario et al., 2021; O’Connor and Zeffiro, 2019; Smitha et al., 2017) (such as patients in vegetative state, coma, etc.). As rs-fMRI does not require overt task engagement, variability related to task performance, effort, or comprehension is minimized (Rosazza and Minati, 2011), thereby improving comparability across subjects and experimental settings (Mulders et al., 2015). This approach enabled the detection of reproducible RSFC patterns and resting-state networks (RSN), which comprise brain areas that exhibit strong RSFC (Beckmann et al., 2005; Biswal et al., 2010; Fransson, 2005; Salvador et al., 2005; van de Ven et al., 2004; van den Heuvel and Hulshoff Pol, 2010). RSNs demonstrate consistency in their large-scale architecture across individuals (Biswal et al., 2010) and support a wide range of cognitive skills and daily-life functioning (Hausman et al., 2020; Kraft et al., 2019). Typical examples include the default mode (DMN), sensorimotor (SMN), central executive (CEN), attentional (AN), language (LN), visual (VN), and salience network (SN) (Manan et al., 2020; Smith et al., 2009; Smitha et al., 2017; van den Heuvel and Hulshoff Pol, 2010; Yeo et al., 2011).

Because RSNs reflect the brain’s task-independent, intrinsic functional organization, they offer a valuable framework for understanding how baseline network dynamics relate to cognitive, emotional, and behavioral functioning. Alterations within or between RSNs have been linked to symptoms and functional impairments across various neurological, neuropsychological, and psychiatric disorders (Rosazza and Minati, 2011; Smitha et al., 2017; Soldozy et al., 2020; van den Heuvel and Hulshoff Pol, 2010). Therefore, modeling RSNs can advance our understanding of both typical psychological processes and diverse forms of psychopathology.

### 1.1 RSNs in Neurology

A substantial body of research documents alterations in functional connectivity across a range of neurological and neurodegenerative diseases (Guggisberg et al., 2008; Harris et al., 2014; Manan et al., 2020; Rigolo et al., 2011; van den Heuvel et al., 2008; Zhou et al., 2012). These alterations frequently involve large-scale networks, such as the DMN (van den Heuvel and Hulshoff Pol, 2010; Wang et al., 2024; Zhou et al., 2012). In Alzheimer’s disease, for instance, pathology may initially affect the DMN in early stages and subsequently propagate into other networks depending on their connectivity to the disease epicenter, with highly connected networks being most vulnerable (Zhou et al., 2012). In other conditions, with primary white matter pathology, for instance in multiple sclerosis, reductions in structural connectivity directly affect FC within sensory-motor networks (van den Heuvel and Hulshoff Pol, 2010). Together, these findings illustrate how RSNs can provide insights into disease progression and network vulnerability, thereby informing disease models with potential relevance to support treatment.

In neurological conditions with focal brain damage, such as lesions or tumors, FC alterations can serve as markers that can guide clinical decision making. For instance, in patients with brain tumors, RSNs may be disrupted by the presence of the tumor (Guggisberg et al., 2008; Manan et al., 2020) its progression (Manan et al., 2020), and surrounding necrotic or infiltrated tissue that no longer participates in functional interactions (Guggisberg et al., 2008; Manan et al., 2020). The extent and pattern of disruption depend on tumor type, grade, location, and growth dynamics (Manan et al., 2020). Although disruptions are typically most pronounced in networks ipsilateral to the tumor (Manan et al., 2020), they can also extend contralaterally (Hadjiabadi et al., 2018), and manifest as reduced intra- and inter-hemispheric connectivity across multiple networks (e.g. DMN, SMN, LN, VN). Importantly, such alterations have been associated with impaired neuropsychological performance (Manan et al., 2020).

Beyond characterizing network disruption, RSN integrity can be informative to clinicians regarding rehabilitation and treatment strategies in patients with focal brain damage. In brain tumor patients, tumor-infiltrated areas may remain part of RSNs important for high-order cognition, such that their resection can cause significant deficits (Guggisberg et al., 2008; Mandal et al., 2024). Conversely, regions with low local functional connectivity can often be removed with minimal functional loss (Guggisberg et al., 2008). More broadly, focal lesions can affect patients’ physical, emotional, cognitive, and social functioning, underscoring the relevance of network-related markers for quality of life (Manan et al., 2020; Warren et al., 2014). For example, higher RSN connectivity has been shown to predict better patient-reported outcomes in individuals with gliomas (Heinzel et al., 2023). Although structural damage in focal lesions is typically spatially localized, RSNs can reveal how the broader network topology is reconfigured following the lesion (Guggisberg et al., 2008; Mandal et al., 2024); and altered interconnected systems can explain cognitive and behavioral impairment (Warren et al., 2014). Together, these findings highlight the importance of monitoring RSNs for both research and clinically informative purposes, as they provide valuable information for predicting clinical and treatment-related outcomes (Liu et al., 2009; Manan et al., 2020; O’Connor and Zeffiro, 2019; Smitha et al., 2017).

### 1.2 Limitations of fMRI for Clinical RSN Mapping

Despite the strengths of rs-fMRI for mapping RSNs, and monitoring network reorganization, its widespread integration into routine clinical practice remains limited (Karambelkar et al., 2022; O’Connor and Zeffiro, 2019; Smitha et al., 2017; Sorensen, 2006). Several practical constraints contribute to this gap. First, fMRI is highly sensitive to motion of the participant which can substantially degrade signal quality (Rosazza and Minati, 2011), and requiring patients to be motionless inside a confined, noisy environment can be particularly challenging for many clinical populations (Manuweera et al., 2025) Second, fMRI acquisition is restricted to hospital settings and cannot be readily implemented in a home-based (or a more ecologically valid) setting (Abdalmalak et al., 2022). Moreover, longitudinal assessment is often required to track functional reorganization and evaluate treatment effects. However, frequently repeated fMRI sessions are often impractical due to limited scanner availability (Karambelkar et al., 2022). clinical scheduling constraints (Karambelkar et al., 2022; Manuweera et al., 2025), safety considerations related to strong magnetic fields (Manuweera et al., 2025; Rigolo et al., 2011) and the substantial financial burden of multiple sessions (Etkin and Mathalon, 2024; Hu et al., 2020; Manuweera et al., 2025). Finally, although resting-state paradigms are straightforward to administer, fMRI data acquisition and analysis require specialized expertise, including operating complex hardware and software (O’Connor and Zeffiro, 2019; Rigolo et al., 2011).

Given these barriers, there is a clear need for complementary tools that allow reliable longitudinal RSN monitoring in clinical contexts. A promising alternative technique is functional near-infrared spectroscopy (fNIRS), which quantifies oxygenated (HbO) and deoxygenated hemoglobin (HbR). Although fNIRS is restricted to measuring hemodynamic activity in the superficial layers of the cortex and has lower spatial resolution relative to fMRI (Cui et al., 2011; Fantini et al., 2018; Quaresima and Ferrari, 2019), its coverage may nevertheless be sufficient to reliably assess FC among cortical areas. Furthermore, fNIRS provides several practical advantages: It is easier to use (Fantini et al., 2018; Kohl et al., 2020), requires little set-up time (Kohl et al., 2020), it is safe, suitable for virtually all populations, portable, without acoustic noise, and tolerant to motion (Fantini et al., 2018; Kohl et al., 2020; Pinti et al., 2020). Most importantly, fNIRS is markedly more cost-effective than fMRI (Fantini et al., 2018; Kohl et al., 2020; Pinti et al., 2020). Together, these features position fNIRS as a strong candidate for group-level longitudinal monitoring of connectome dynamics in neurological patients.

### 1.3 fNIRS for RSN Monitoring

While numerous studies have demonstrated the ability of fNIRS to estimate RSFC (Einalou et al., 2017; Lu et al., 2010; Mesquita et al., 2010; Niu et al., 2013; Zhang et al., 2010; Zhang, 2010), the extent to which it can reliably identify large-scale RSFC patterns and RSNs corresponding to those derived from fMRI remains unclear (Aihara et al., 2020; Duan et al., 2012; Sasai et al., 2012). For example, Duan et al. (2012) used simultaneous fMRI-fNIRS recordings to examine RSFC of the motor cortex. The authors found higher correspondence in RSFC patterns between fMRI and fNIRS when using the HbO signal than with HbR, and comparable local efficiency and characteristic path length in binarized graphs (Duan et al., 2012). While HbR is generally more closely related to neural activity than HbO (Montero-Hernandez et al., 2018), it suffers from lower signal-to-noise ratio (Montero-Hernandez et al., 2018; Pinti et al., 2020), and it remains unclear which fNIRS signal better reflects the BOLD response (Murata, 2002; Steinbrink et al., 2006; Strangman et al., 2002). With this context in mind, Duan et al. (2012) did not use short-channel regression to remove the superficial scalp noise from the fNIRS data, which is a critical preprocessing step. Therefore, it is uncertain whether the higher correspondence observed for HbO reflects true alignment with BOLD, or whether the HbR signal was confounded by uncorrected noise. In another study, Aihara et al. (2020) compared FC matrices of frontal and parietal regions from the two modalities and reported only moderate agreement. Overall, previous work has largely focused on limited regions, small cohorts, or single analytical levels (e.g. edgewise correlations or binarized network metrics), and has often omitted critical preprocessing steps, leaving unresolved questions about the true cross-modal correspondence of fNIRS and fMRI RSFC patterns and whether fNIRS can reliably monitor whole-brain RSNs or support clinical applications requiring higher-order network metrics for informed decision making.

To address these gaps, we conducted a comprehensive, multi-level comparison of resting-state connectivity derived from fMRI and fNIRS. Functional connectivity was estimated using both bivariate and partial correlations to assess the impact of methodological choices on cross-modal correspondence. Similarities and differences were examined across multiple scales, including edge-, nodal-, graph-, and modular-level organization, incorporating higher-order metrics such as nodal strength, assortativity, local efficiency, path length, and betweenness centrality. Applying this framework with near whole-head fNIRS coverage, we provide the first systematic, multi-scale evaluation of fNIRS–fMRI correspondence, clarifying where fNIRS converges with fMRI, where discrepancies persist, and which aspects of network organization are robust across modalities. These findings establish a critical foundation for evaluating the translational utility of fNIRS in clinical research and for guiding methodological choices in future cross-modal connectivity studies.

## 2 Methodology

### 2.1 Participants

The current study analyzed data from two independent cohorts. fNIRS data was obtained from a dataset that is currently being prepared for publication; the analyses reported here were conducted on the data available at the time (*N =* 49) and therefore does not include the final full sample. Subjects were excluded if more than 30% of either short or long channels had a scalp coupling index (SCI) below 0.7 (*N* = 17), or if resting-state fNIRS data were missing (*N* = 1). The final sample included 31 healthy participants (mean age = 28.25 years; SD = 11.11; 61.29% female). During acquisition, subjects completed a 5-min resting-state run and were instructed to clear their mind while fixating on a cross at the center of the screen.

For resting-state fMRI, data were obtained from the Human Connectome Project (HCP) (Van Essen et al., 2012) and were individually matched to the fNIRS cohort for age and gender. Each HCP participant completed a 15-min resting-state scan while fixating on a crosshair. All procedures were conducted in accordance with the Declaration of Helsinki. The fNIRS data were collected as part of a study approved by the local institutional ethics committee, and all participants provided written informed consent prior to participation. Resting-state fMRI data were obtained from the HCP, which received approval from the Washington University Institutional Review Board, and all HCP participants provided written informed consent for data sharing and secondary analysis.

### 2.2 fNIRS & fMRI Acquisition and Preprocessing

Hemodynamic signals were recorded with two cascading NIRSport2 devices (NIRx Medical Technologies, Berlin) and Aurora v2021.1 software. The fNIRS setup included 32 LED sources, 28 silicon photodiode detectors, with 32 short distance detectors 8mm from each source (134 channels in total; λ_1_ = 760 nm, λ_2_ = 850 nm). Optode placement was guided using the fOLD toolbox (Zimeo Morais et al., 2018) and arranged according to the international 10-10 system. Processing of fNIRS data was performed with Satori 2.2.4 (Brain Innovation, Maastricht, Netherlands) and Python v3.11.9. The first and last 20 seconds of the resting-state data were trimmed to reach steady state (Duan et al., 2012; Lu et al., 2010; Zhang, 2010). Raw signals were converted to optical density and subsequently to hemoglobin concentrations using the Beer-Lambert law. Channels with SCI below 0.7 were rejected and motion artifacts were corrected (spike removal combined with temporal derivative distribution repair) (Khan et al., 2024). Short-channel regression was performed using the general linear model based on the highest correlated short-channel. Finally, the data were bandpass filtered (0.01–0.1 Hz) (Liu et al., 2023; Notte et al., 2024; Smitha et al., 2017; Zhang et al., 2010) to preserve resting-state fluctuations and remove physiological noise (Duan et al., 2012; Liu et al., 2023; Lu et al., 2010; Zhang et al., 2010), linear trends were removed and the resulting time-series was *z*-normalized.

Structural and functional MRI data were acquired according to the standard HCP protocols on a 3T Siemens scanner (Feinberg et al., 2010; Milchenko and Marcus, 2013; Moeller et al., 2010; Setsompop et al., 2012; Sotiropoulos et al., 2013; Van Essen et al., 2012; Xu et al., 2012). High-resolution T1-weighted (MPRAGE; TR = 2400 ms; TE = 2.14 ms) structural images were obtained with at 0.7 mm isotropic resolution. Functional BOLD fMRI was acquired with a multiband EPI sequence (2.0 mm isotropic, 72 slices, TR = 720 ms, TE = 33.1 ms, flip angle 52°, multiband factor = 8) with both right-to-left and left-to-right phase encoding. Preprocessing followed the HCP pipeline, including gradient distortion and motion correction using SBRef image, EPI distortion correction using opposite phase-encoding scans and co-registration; rs-fMRI data were further cleaned using HCP’s ICA-FIX procedure, which uses MELODIC ICA (Fischl, 2012; Glasser et al., 2013; Jenkinson et al., 2012, 2002). For the present analysis, the recommended HCP preprocessed data were used from the first session with left-to-right encoding (e.g. rfMRI_REST1_LR_hp2000_clean_rclean_tclean.nii.gz).

### 2.3 Channel-to-ROI mapping

For each of the fNIRS channels (n=102), MNI coordinates were determined using the fOLD toolbox (Zimeo Morais et al., 2018), which maps channels to standard anatomical locations based on validated parcellation methods and the 10-10 system. From the original fNIRS acquisition, signals of 20 channels were excluded because fOLD could not reliably provide MNI coordinates for these channel pairs in our montage. For the remaining 82 channels, spherical ROIs (0.5 cm radius) were created with FSL (Jenkinson et al., 2012) at their respective MNI coordinates. Due to the limited spatial resolution and variability in fNIRS optode placement, these ROIs provide an approximation rather than precise mapping of cortical locations. Then the average preprocessed BOLD signal across all voxels within each ROI sphere was extracted for each HCP subject (using Nibabel 5.3.2 and Nilearn 0.11.1), resulting in corresponding timeseries between fNIRS and fMRI ROIs (e.g. channel S1D1 to ROI1).

### 2.4 Resting State Functional Connectivity Matrices

Separate RSFC matrices were created based on fMRI, HbO and HbR fNIRS signals. For each subject, a correlation matrix was computed by correlating the time-series of each ROI with every other ROI. Both bivariate and partial correlations were calculated. For each subject (*n* = 31), modality-specific signal (*n* = 3) and correlation type (*n* = 2), an 82 × 82 correlation matrix was created.

To obtain the group-level RSFC per modality, each subject-level correlation matrix was first transformed using Fisher’s *z*, which converts correlation coefficients to an approximately normally distributed scale (Silver and Dunlap, 1987). The *z*-transformed matrices were then averaged across subjects, yielding six group-level correlation matrices (i.e. partial vs. bivariate correlations × 3 signal types).

### 2.5 Similarity Between Modalities

To evaluate the similarity of connectivity patterns between fNIRS and fMRI, both subject-level and group-level analyses were conducted.

First, based on individual matrices, independent samples *t-*tests, with Benjamini-Hochberg false discovery rate (BH FDR) correction, were performed to assess edgewise differences in functional connectivity between modalities across subjects. To complement this, the two one-sided *t*-tests (TOST) procedure (Schuirmann, 1981) (with BH FDR corrections) was applied to evaluate statistical equivalence of edges between modalities across a range of equivalence bounds. Performing TOST allows to determine whether fNIRS and fMRI RSFC edge-values are not merely non-significantly different but can be considered interchangeable within a specified correlation difference. Prior to the edgewise *t-*tests and TOSTs, Shapiro-Wilk tests were performed for each edge to verify that the normality assumption was satisfied.

Second, based on the group-averaged correlation matrices, correspondence between fNIRS and fMRI RSFC patterns was assessed using Pearson correlation, evaluating whether the overall structure of functional connections is preserved across modalities. To account for the non-independence of ROI pairs, significance was determined via permutation testing (*N* = 100,000), in which the row and column labels of the fMRI matrix were randomly shuffled while keeping the fNIRS matrix fixed. This generated a null distribution of correlation values expected by chance, and the *p*-value was calculated as the proportion of permuted correlations with absolute values equal to or exceeding the observed correlation, providing a robust estimate of cross-modal similarity.

Third, nodal graph metrics were computed, to assess whether fNIRS and fMRI capture comparable patterns of regional and large-scale network organization beyond individual connections (i.e. net nodal strength, normalized nodal strength (Rubinov and Sporns, 2011), assortativity, local efficiency, path length, and betweenness centrality). Whereas edge-level analyses focus on specific pairwise relationships, nodal metrics summarize how each region participates in the network as a whole, and modular analyses test whether both modalities recover similar functional subnetworks. On the subject-level, nodal metric differences between modalities were analyzed with independent samples *t-*tests (with FDR BH correction). On the group-level, for each node metric, the spatial relationship across modalities was explored with correlational analysis.

Finally, modularity was estimated for each modality-specific group-averaged matrix using the Louvain algorithm, with stability being ensured through repeated optimization and selection of consistent community assignments. Spatial overlap of modules between fMRI and fNIRS was quantified using Jaccard similarity and statistical significance was assessed via permutation testing.

All abovementioned cross-modal comparisons were conducted for fMRI vs. both fNIRS HbO and HbR, using both bivariate and partial correlation matrices. Python v3.11.9 was used for all analyses, unless stated otherwise; for more details on the methodology and implementation of analyses see Appendix A.

## 3 Results

The group-RSFC matrices of fMRI and fNIRS (HbO/HbR), computed using bivariate and partial correlations, are shown in Fig. 1. These matrices are Fisher *z*-transformed, normalized by the global maximum, and reordered according to their stable communities (see Appendix A, Section 13.3 and 13.4 for details). The corresponding raw group-RSFC matrices, ordered by the original channel/ROI arrangement, are shown in Fig. B1, with their edge-value distributions in Fig. B.2.

**Figure 1.**
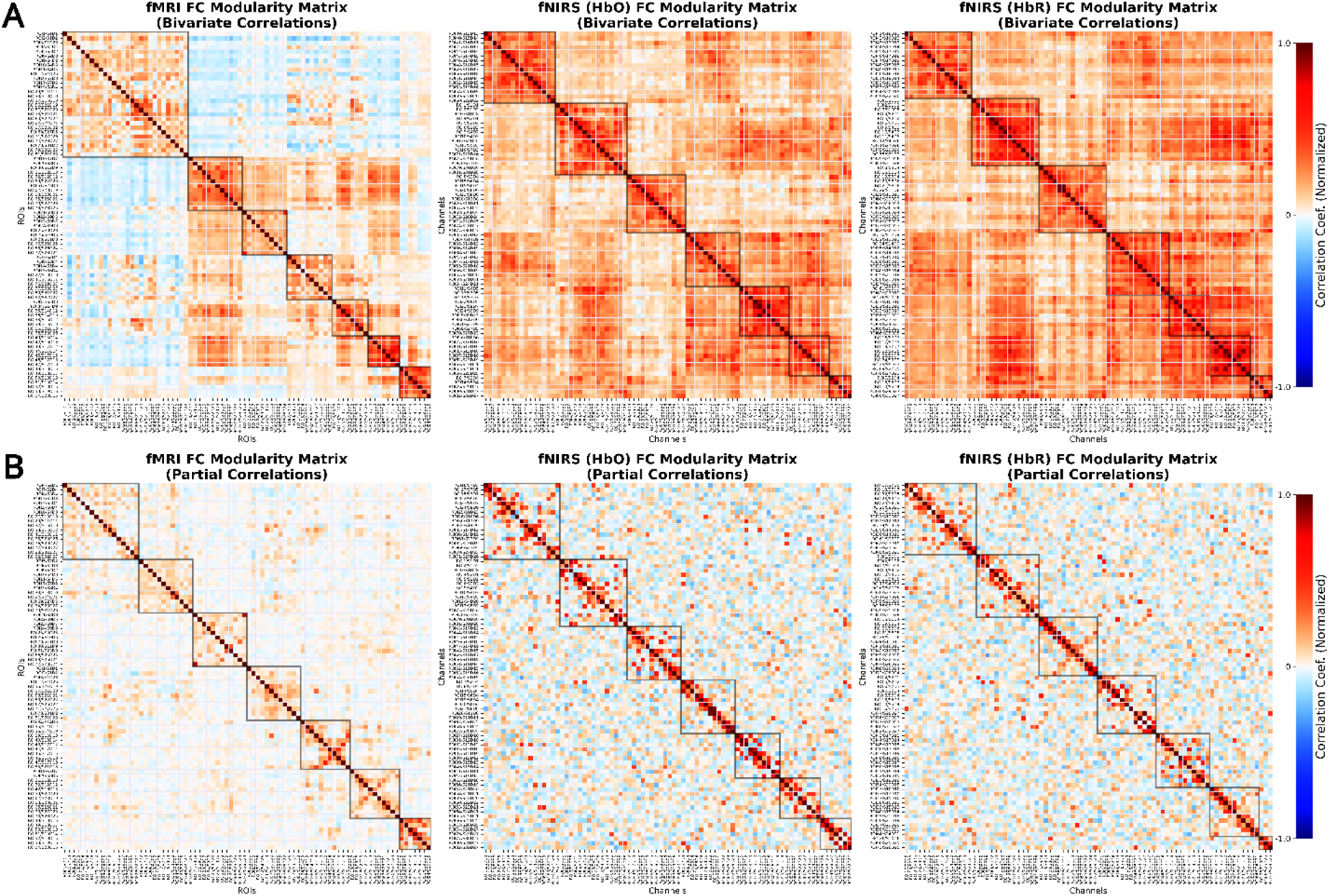
Group-level RSFC matrices sorted by modularity. Panel (A) represents RSFC matrices derived from bivariate and panel (B) from partial correlations. Matrices represent the Fisher *z*-transformed, group-averaged correlations, normalized by the global maximum across modalities. Louvain modularity was computed across a range of gamma values to identify the gamma producing seven-stable communities. Because the Louvain algorithm is greedy, different runs produce different community assignments. To capture this variability, we performed 1000 iterations (Rubinov and Sporns, 2011). Then with a consensus clustering algorithm a single, stable community assignment was generated (Bassett et al., 2013; Sporns and Betzel, 2016). Channels/ROIs were reordered according to the consensus communities and subsequently by community size. Square boxes outline the sets of ROIs belonging to each community.

### 3.1 Differences & Similarities Between Modalities

#### 3.1.1 Edgewise t-tests

Shapiro-Wilk tests verified that most edge vectors were normally distributed for both bivariate and partial correlation estimates of RSFC (resp. 78.38%/89.40% of fMRI, 92.92%/93.74% of fNIRS-HbO, 90.15%/94.58% fNIRS-HbR edges).

Independent samples *t*-tests revealed that 50.29% of edges significantly differed between fMRI and fNIRS-HbO and 61.43% of edges differed between fMRI and fNIRS-HbR. Based on partial correlation matrices, only 2.14% for fMRI vs fNIRS-HbO and 1.26% for fMRI vs fNIRS-HbR of edges were significantly different, respectively (Fig. B.3).

#### 3.1.2 Edgewise TOST

Based on bivariate correlations, at least 50% of the fMRI edges were equivalent to fNIRS edges when the difference in edge-values was larger than 0.207 for HbO and 0.245 for HbR. With partial correlation, 50% of the fMRI edges were equivalent to fNIRS edge-values when the difference in edge-values was at least 0.111 for HbO and 0.112 for HbR (Fig. B.4). These thresholds mark the minimum correlation difference levels at which half of the connectome demonstrates cross-modality agreement.

#### 3.1.3 Matrix-level correspondence of RSFC patterns

To quantify the similarity between fMRI and fNIRS overall RSFC patterns, Pearson correlations were computed between group-averaged connectivity matrices, with statistical significance assessed using permutation testing (100,000 permutations). Using bivariate correlations, fMRI RSFC showed moderate and significant correspondence with both fNIRS chromophores (HbO: *r* = 0.393, *P* < 10^-5^; HbR: *r* = 0.353, *P* < 10^-5^). Comparable results were observed for partial correlation-based RSFC estimates, with significant matrix-level similarity for both HbO (*r* = 0.379, *P* < 10^-5^) and HbR (*r* = 0.385, *P* < 10^-5^); indicating a shared large scale connectivity organization between fMRI and fNIRS RSFC, which was robust across correlation types and hemoglobin signals.

### 3.2 Graph Metrics

#### 3.2.1 Similarity of nodal metrics

Graph metrics were calculated both on subject- and group-level matrices (separately for bivariate and partial correlations). For each node, a vector containing the metric values across subjects was constructed separately for fMRI, fNIRS-HbO, and fNIRS-HbR. These vectors were then compared pairwise using independent samples *t*-tests, with BH FDR correction. Table 1 reports the percentage of nodes showing significant differences between modalities for net nodal strength, normalized nodal strength, assortativity, nodal efficiency, and nodal path length. Nodewise differences for betweenness centrality were not assessed, because the dense weighted functional connectivity graphs yielded trivial (zero) betweenness centrality values for most nodes across subjects, reflecting the dominance of the shortest direct paths in fully connected networks.

**Table 1.** Percentage of nodewise differences between modalities.

| Graph metric | fMRI vs fNIRS (HbO) |  | fMRI vs fNIRS (HbR) |  |
| --- | --- | --- | --- | --- |
|  | Bivariate | Partial | Bivariate | Partial |
| Net Strength | 79.27% | 37.80% | 89.02% | 37.80% |
| Normalized Strength | 0.00% | 25.61% | 0.00% | 19.51% |
| Assortativity | 23.17% | 14.63% | 29.27% | 24.39% |
| Local Efficiency | 47.56% | 100.00% | 80.49% | 100.00% |
| Path Length | 39.02% | 100.00% | 65.85% | 100.00% |
| Percentage of nodes showing significant differences (BH FDR corrected) in subject-level nodal centrality metrics between modalities. |  |  |  |  |

Table 2 reports the similarity in the pattern of nodal graph metrics derived from group-average matrices between modalities. For each graph metric, a vector was formed containing the nodal values across all ROIs for fMRI, and this was correlated with the corresponding nodal vector of fNIRS-HbO and fNIRS-HbR. Because each ROI in fMRI maps directly to the same ROI in fNIRS this allowed to assess the similarity between modalities in network organization of graph properties. The choice of correlation (Pearson’s *r* or Spearman’s *rho*) was determined by Shapiro-Wilk tests on the nodal distributions. Table 2 reports these inter-modality correlations for net nodal strength, normalized nodal strength, assortativity, nodal efficiency, and nodal path length. Similarity in the distribution of nodal betweenness centrality was not assessed for group-averaged matrices, as betweenness centrality produced trivial (zero) values for all nodes in the dense weighted graph.

**Table 2.** Similarity in the distribution of nodal metrics between modalities.

|  | fMRI vs fNIRS (HbO) |  | fMRI vs fNIRS (HbR) |  |
| --- | --- | --- | --- | --- |
|  | Correlation Statistic | *P*-value | Correlation Statistic | *P*-value |
| **Bivariate Correlations** |  |  |  |  |
| Net Strength | $\rho = 0.179$ | 0.1068 | $\rho = -0.069$ | 0.5393 |
| Normalized Strength | **$\rho = 0.510$** | **<0.0001** | **$\rho = 0.334$** | **0.0022** |
| Assortativity | $\rho = 0.213$ | 0.0548 | **$\rho = 0.421$** | **0.0001** |
| Local Efficiency | **$\rho = 0.492$** | **<0.0001** | **$\rho = 0.255$** | **0.0208** |
| Path Length | **$\rho = 0.432$** | **<0.0001** | $\rho = 0.197$ | 0.0752 |
| **Partial Correlations** |  |  |  |  |
| Net Strength | **$r = 0.224$** | **0.0282** | $r = 0.049$ | 0.6629 |
| Normalized Strength | $\rho = 0.083$ | 0.4591 | $\rho = -0.048$ | 0.6674 |
| Assortativity | $\rho = -0.080$ | 0.4758 | $\rho = 0.073$ | 0.5149 |
| Local Efficiency | $r = 0.0469$ | 0.6758 | $r = -0.0824$ | 0.4616 |
| Path Length | $r = 0.1080$ | 0.3340 | $r = -0.0352$ | 0.7534 |

Graph metrics were calculated based on group-RSFC matrices.
Significance at $P < 0.05$ marked in bold.

#### 3.2.2 Modularity

Stable community structure was identified for each modality on the group average matrices using the Louvain algorithm across a range of resolution *γ* values. Based on visual inspection and qualitative comparison with established canonical RSNs (Yeo et al., 2011) together with considerations of the spatial resolution and cortical coverage of the 82-channel fNIRS montage, the seven-community solution was selected as the most biologically plausible. Modularity was consistently higher for fMRI than fNIRS across both bivariate and partial correlation approaches to estimate RSFC. From bivariate correlations, modularity was *Q* = 0.3961 for fMRI, *Q* = 0.1650 for fNIRS-HbO and *Q* = 0.1474 for fNIRS-HbR. For partial correlation matrices, modularity remained higher for fMRI (*Q* = 0.5215), compared to fNIRS-HbO (*Q* = 0.3578) and fNIRS-HbR (*Q* = 0.3501). Community assignments are shown in Fig. 1, for both bivariate and partial correlation matrices.

The correspondence between fMRI and fNIRS community structure was quantified using the Jaccard index with statistical significance assessed with permutation testing. For each fMRI community, the fNIRS community with the highest overlap was identified as its best match. For bivariate correlations, the mean of these maximum overlaps across the seven fMRI communities was ∼0.35 across fNIRS chromophores. Several fMRI communities showed statistically significant similarity with fNIRS modules, including the 1^st^, 2^nd^, 3^rd^, 6^th^, and 7^th^ with HbO signal, and 1^st^, 2^nd^, 3^rd^, 5^th^, 6^th^, and 7^th^ with HbR (see Table 3). From partial correlations, the mean maximum overlap was 0.305 for HbO and 0.361 for HbR. The following fMRI communities showed significant overlap with fNIRS-HbO: 1^st^, 2^nd^, 4^th^, 5^th^, 6^th^, and 7^th^; and fNIRS-HbR: 1^st^, 3^rd^, 5^th^, 6^th^, and 7^th^. Individual community overlaps are summarized in Table 3.

**Table 3.**
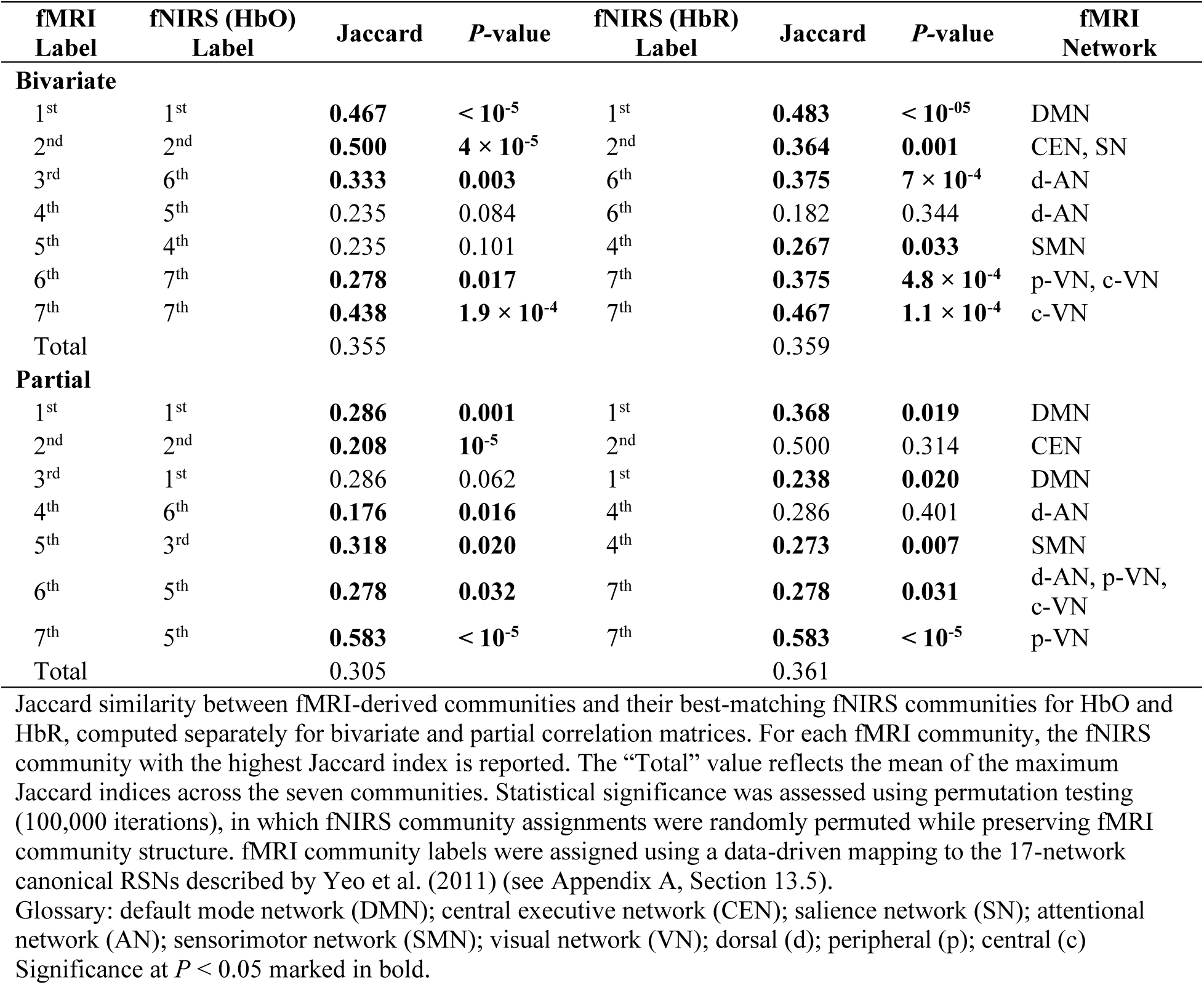
Jaccard similarity analysis on fMRI-fNIRS subnetwork overlap.

| fMRI Label | fNIRS (HbO) Label | Jaccard | *P*-value | fNIRS (HbR) Label | Jaccard | *P*-value | fMRI Network |
| --- | --- | --- | --- | --- | --- | --- | --- |
| **Bivariate** |  |  |  |  |  |  |  |
| 1<sup>st</sup> | 1<sup>st</sup> | **0.467** | **< 10<sup>-5</sup>** | 1<sup>st</sup> | **0.483** | **< 10<sup>-05</sup>** | DMN |
| 2<sup>nd</sup> | 2<sup>nd</sup> | **0.500** | **4 × 10<sup>-5</sup>** | 2<sup>nd</sup> | **0.364** | **0.001** | CEN, SN |
| 3<sup>rd</sup> | 6<sup>th</sup> | **0.333** | **0.003** | 6<sup>th</sup> | **0.375** | **7 × 10<sup>-4</sup>** | d-AN |
| 4<sup>th</sup> | 5<sup>th</sup> | 0.235 | 0.084 | 6<sup>th</sup> | 0.182 | 0.344 | d-AN |
| 5<sup>th</sup> | 4<sup>th</sup> | 0.235 | 0.101 | 4<sup>th</sup> | **0.267** | **0.033** | SMN |
| 6<sup>th</sup> | 7<sup>th</sup> | **0.278** | **0.017** | 7<sup>th</sup> | **0.375** | **4.8 × 10<sup>-4</sup>** | p-VN, c-VN |
| 7<sup>th</sup> | 7<sup>th</sup> | **0.438** | **1.9 × 10<sup>-4</sup>** | 7<sup>th</sup> | **0.467** | **1.1 × 10<sup>-4</sup>** | c-VN |
| Total |  | 0.355 |  |  | 0.359 |  |  |
| **Partial** |  |  |  |  |  |  |  |
| 1<sup>st</sup> | 1<sup>st</sup> | **0.286** | **0.001** | 1<sup>st</sup> | **0.368** | **0.019** | DMN |
| 2<sup>nd</sup> | 2<sup>nd</sup> | **0.208** | **10<sup>-5</sup>** | 2<sup>nd</sup> | 0.500 | 0.314 | CEN |
| 3<sup>rd</sup> | 1<sup>st</sup> | 0.286 | 0.062 | 1<sup>st</sup> | **0.238** | **0.020** | DMN |
| 4<sup>th</sup> | 6<sup>th</sup> | **0.176** | **0.016** | 4<sup>th</sup> | 0.286 | 0.401 | d-AN |
| 5<sup>th</sup> | 3<sup>rd</sup> | **0.318** | **0.020** | 4<sup>th</sup> | **0.273** | **0.007** | SMN |
| 6<sup>th</sup> | 5<sup>th</sup> | **0.278** | **0.032** | 7<sup>th</sup> | **0.278** | **0.031** | d-AN, p-VN, c-VN |
| 7<sup>th</sup> | 5<sup>th</sup> | **0.583** | **< 10<sup>-5</sup>** | 7<sup>th</sup> | **0.583** | **< 10<sup>-5</sup>** | p-VN |
| Total |  | 0.305 |  |  | 0.361 |  |  |

Jaccard similarity between fMRI-derived communities and their best-matching fNIRS communities for HbO and HbR, computed separately for bivariate and partial correlation matrices. For each fMRI community, the fNIRS community with the highest Jaccard index is reported. The “Total” value reflects the mean of the maximum Jaccard indices across the seven communities. Statistical significance was assessed using permutation testing (100,000 iterations), in which fNIRS community assignments were randomly permuted while preserving fMRI community structure. fMRI community labels were assigned using a data-driven mapping to the 17-network canonical RSNs described by Yeo et al. (2011) (see Appendix A, Section 13.5).
Significance at $P < 0.05$ marked in bold.

These overlaps are illustrated in Fig. 2 using BrainNet Viewer (Xia et al., 2013) where fNIRS-HbO and fNIRS-HbR community assignments are shown after restricting the nodes that did not spatially overlap with their corresponding fMRI community, facilitating direct visual comparison of cross-modal community alignment.

**Figure 2.**
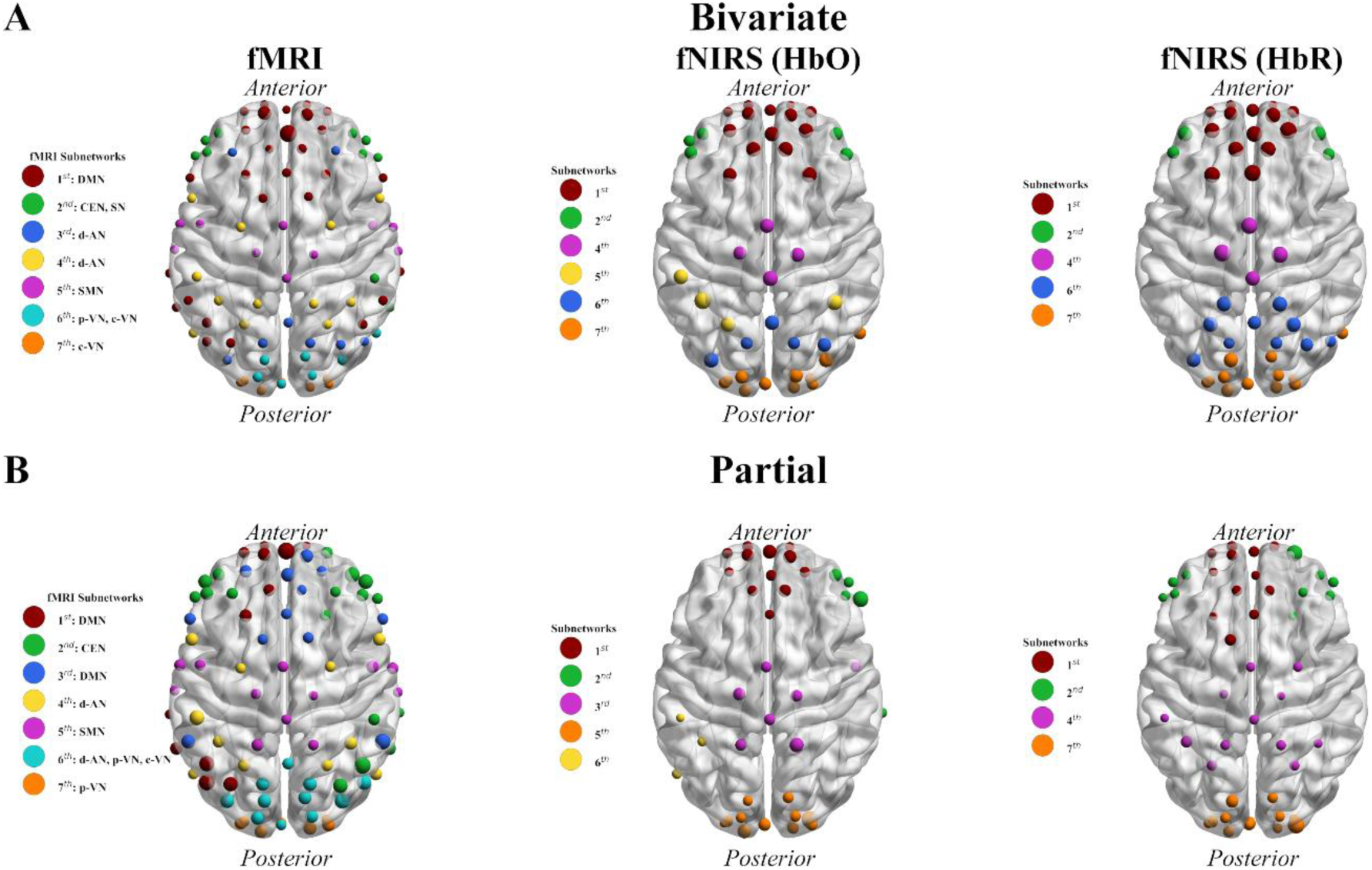
Stable community structure of the group-average correlation matrices. Panel (A) represents the seven stable communities derived from bivariate matrices and (B) those derived from partial matrices. Nodes of fMRI communities are colored according to their assigned network labels. Network labels are based on the 17-canonical resting-state networks from Yeo et al. (2011) (see Appendix A). In fNIRS HbO and HbR communities, only the nodes that spatially overlap with each corresponding fMRI community are displayed. Glossary: default mode network (DMN); central executive network (CEN); salience network (SN); attentional network (AN); sensorimotor network (SMN); visual network (VN); dorsal (d); peripheral (p); central (c).

## 4 Discussion

The present study evaluated the extent to which fNIRS can serve as a clinically relevant tool for RSFC estimation by direct comparison with fMRI. Unlike prior work, which was limited to specific regions or thresholded networks, a comprehensive multi-scale assessment across edge-, node-, graph-, and module-level analyses was performed using weighted bivariate and partial correlation matrices. This allowed us to capture the full range of connectivity, and avoided arbitrary thresholds (van Wijk et al., 2010), preserving subtle network features. At the group-level, fNIRS reliably captured large-scale RSFC patterns, graph-metric distributions and modular architecture observed with fMRI, whereas cross-modal correspondence diminished at the subject-level. Together, these results provide the first quantitative demonstration of fNIRS-fMRI correspondence across multiple analytical scales. These findings support the translational potential of fNIRS for group-level RSFC mapping and provide a framework for evaluating methodological choices that shape cross-modal correspondence.

### 4.1 Edgewise and global RSFC correspondence

With connectivity matrices at the subject-level, partial correlations showed fewer edgewise differences between modalities than bivariate correlations. To better interpret these findings, the TOST procedure was applied, which allows to formally assess whether edges can be considered statistically equivalent across modalities. Unlike traditional *t*-tests, which only indicate whether differences are statistically significant, TOST tests whether the observed differences fall within a predefined, practically negligible range. Because no prior work has established equivalence bounds for cross-modal RSFC, we explored a range of bounds (Δ*r* ≈ 0.01–0.4) to determine the interval over which edges could be considered equivalent. Using this approach, we found that half of the edges reached equivalence at very narrow bounds (∼0.10) for partial correlations, whereas bivariate correlations required considerably wider bounds (∼0.25) to achieve a comparable proportion of equivalent edges.

These results suggest that partial correlations reduce modality-specific differences and emphasize more direct, consistent connectivity patterns across fMRI and fNIRS. In contrast, bivariate correlations may remain inflated by shared variance such as physiological noise or global superficial signal. Nonetheless, while partialling out the shared variance reduces false positives, it may also remove true functional interactions (Fornito et al., 2016a), driving many edge weights towards zero. As a result, the increased inter-modal correspondence observed with partial correlations likely reflects a combination of genuine overlap in direct structurally constrained connectivity (Fornito et al., 2016a; Varoquaux and Craddock, 2013), increased network sparsity and a global reduction in edge weights.

At the group-level, the similarity between fMRI and fNIRS patterns was quantified by correlating their average connectivity matrices and assessed for significance with permutation testing to account for the non-independence of ROI pairs. This analysis revealed a significantly moderate cross-modal correspondence (∼0.35–0.39) between fMRI and fNIRS matrices for both correlation approaches. These results indicate that the overall topology of FC (i.e. the relative pattern of which ROI pairs show stronger or weaker coupling) is broadly preserved across modalities. The observed magnitude of the similarity is consistent with previous work (Aihara et al., 2020), though direct comparison is limited due to methodological differences across studies.

### 4.2 Cross-modal graph metric correspondence

To assess whether fNIRS captures meaningful functional organization beyond individual edges or global RSFC patterns, a range of graph-theoretical metrics were examined. Nodewise differences indicate how individual nodes vary across modalities, while similarity in overall nodal patterns reflects preservation of the network as a whole. Comparing bivariate and partial correlations further reveals which approach shows greater cross-modal agreement.

Building on this framework, net nodal strength showed limited cross-modal comparability, especially for bivariate correlations, with significant nodewise differences observed in ∼80–90% of nodes and no consistent pattern across nodes. This likely reflects its sensitivity to negative weights, which were more prominent in fMRI bivariate matrices and may amplify inter-modality differences.

In contrast, normalized nodal strength, which downweights negative edges and is more robust in signed networks (Fornito et al., 2016b; Rubinov and Sporns, 2011) showed the strongest cross-modal agreement. Under bivariate correlations, no nodes differed significantly, and nodewise patterns were moderately to strongly correlated across chromophores. Partial correlations modestly increased nodewise differences (≈20–25% of nodes) and largely lost any similarity across nodes. This indicates that metrics accounting for edge sign, combined with bivariate estimation of RSFC, provide more stable and comparable estimates of regional network importance. This supports normalized nodal strength as possibly valuable metric for translational fNIRS-fMRI comparisons.

Assortativity showed minimal nodal differences between modalities (≈15–30% of nodes), indicating relative stability at the regional level. Across nodes, patterns were modest, reaching significance mainly for bivariate correlations with fNIRS-HbR and trending for fNIRS-HbO, suggesting that assortativity is partially preserved across modalities.

Substantial nodewise differences were observed for local efficiency and nodal path length, particularly under partial correlations, where all nodes differed significantly between modalities. Under bivariate correlations, nodewise patterns showed small-to-moderate similarity, especially for HbO, consistent with prior work on thresholded matrices (Duan et al., 2012). This suggests that fNIRS can partially capture broad patterns of local network integration and shortest paths, though absolute nodal estimates remain modality dependent. Interestingly, although signed weighted matrices are generally suboptimal for shortest-path metrics (Fornito et al., 2016b, 2016c; Rubinov and Sporns, 2011; van Wijk et al., 2010), and partial correlations theoretically emphasize direct paths (Fornito et al., 2016b), greater cross-modal similarity in path length was observed for bivariate correlations; indicating that correspondence is driven more by shared dense connectivity than by sparse, direct pathways.

Overall, nodewise differences across modalities were pronounced for most centrality metrics, except for normalized nodal strength and assortativity, which showed moderate cross-modal similarity, especially with bivariate correlations. This supports fNIRS utility for these metrics in clinical populations. In contrast, path- or flow-dependent metrics, such as local efficiency and nodal path length, exhibited greater modality dependence and limited convergence.

### 4.3 Correspondence in subnetworks

At the network level, moderate overlap in modular organization was observed between fMRI and fNIRS, and permutation testing confirmed that this similarity exceeded chance levels. These findings indicate that both modalities partition ROIs into broadly comparable functional modules. In more detail, with the bivariate approach, DMN, VNs, and attentional [i.e. CEN, dorsal-AN (d-AN), and SN; fMRI modules 1^st^–3^rd^, 6^th^–7^th^] networks were consistently identified across modalities for both fNIRS chromophores, demonstrating convergence in subnetwork organization. In fMRI the visual network was differentiated in central and peripheral components. In contrast, this finer-grained segregation was not reliably captured by fNIRS across chromophores, where the visual network emerged as a single community.

Based on partial correlation matrices, fMRI subnetwork partitioning was less robust. The SN was not detected, multiple modules corresponded to the DMN, and the 6^th^ module included the d-AN together with both central and peripheral VNs (c-VN; p-VN). Correspondence with fNIRS was also less consistent across chromophores and more diffuse for visual networks, with c-VN and p-VN again merging into a single module. These findings suggest that while broad modular organization is generally preserved across modalities, bivariate correlations yield more stable and chromophore-consistent overlap, whereas finer-grained distinctions within the visual system are not reliably captured by fNIRS and partial correlation graphs yielded less consistent overlap with well-acknowledged RSNs in fMRI.

Collectively, despite the methodological limitations inherent to the present study, fNIRS can identify key topological RSN properties and reproduce large-scale functional cortical subnetworks, within the limits of its spatial coverage. Importantly, convergence between modalities depends on methodological choices (i.e. partial vs. bivariate matrices, HbO vs HbR), highlighting the need for careful selection and interpretation of analytical approaches when applying fNIRS to clinical network mapping.

### 4.5 Clinical Implications

Although the present study was conducted in healthy participants and does not directly inform clinical decision making, it clarifies the conditions under which fNIRS may be clinically informative for RSN assessment. A central finding is the contrast between the moderate cross-modal convergence observed at the group level (e.g., global RSFC patterns, network organization, community structure) and the pronounced divergence at the individual level (e.g., edge- and node-level measures, with limited exceptions). Group-averaged connectivity matrices showed meaningful similarity between fNIRS and fMRI, indicating that fNIRS captures core aspects of large-scale functional architecture in superficial cortical networks and enables reliable identification of major functional subnetworks. This group-level robustness supports the use of fNIRS for characterizing population-level RSN alterations in clinical cohorts, particularly when MRI is contraindicated or impractical.

In contrast, subject-level analyses revealed substantial edgewise and nodewise discrepancies between modalities. Importantly, such variability is not unique to fNIRS, as rs-fMRI itself exhibits considerable inter-subject variability and only moderate test-retest reliability (O’Connor and Zeffiro, 2019; Specht, 2020). These findings underscore that fNIRS should be applied cautiously in clinical contexts requiring precise subject-specific connectivity estimates, as cross-modal divergence becomes more pronounced for fine-grained network features.

Despite these limitations, fNIRS offers clear clinical advantages in applications that prioritize network-level changes over individual edges or nodes. Its portability, low cost, and short setup time (Fantini et al., 2018; Kohl et al., 2020; Pinti et al., 2020) make it well suited for longitudinal monitoring of RSN alterations, enabling repeated assessments of disease progression, recovery, or treatment response. In conditions such as brain tumors and epilepsy, resting-state connectivity alterations have been linked to functional outcomes, supporting the use of network-based markers for risk stratification (Lang et al., 2014). For example, RSFC measures may help anticipate post-operative deficits (Guggisberg et al., 2008; Mandal et al., 2024) or identify network alterations associated with oncogenesis (Bullmore and Sporns, 2012; Mandal et al., 2020) in brain tumor patients, while in epilepsy fNIRS could facilitate quick identification of patients at risk of recurrent seizures following surgery (Lang et al., 2014).

fNIRS also provides a practical platform for tracking network-level effects of neuromodulatory interventions, where repeated measurements are often required and integration with fMRI may be challenging (Lang et al., 2014). In such contexts, the clinical utility of fNIRS lies in capturing treatment-related changes in large-scale network organization rather than fine-grained connectivity. Moreover, as clinical resting-state studies are often limited by small sample sizes (Herold et al., 2017; Lang et al., 2014; Marek et al., 2022; Noble et al., 2019; van Wijk et al., 2010), the accessibility and scalability of fNIRS offer the opportunity to study larger and more diverse cohorts, improving reproducibility and enabling systematic investigation of population-level network alterations (Herold et al., 2017; Pinti et al., 2020).

In sum, while subject-level comparisons highlight important cross-modal differences, the observed group-level convergence indicates that fNIRS is well positioned to support population-level RSN characterization and longitudinal clinical monitoring, particularly in settings where repeated or accessible neuroimaging is essential.

### 4.6 Strengths, Limitations, and Future Directions

The present study offers several methodological and conceptual strengths that advance fNIRS RSFC research. Notably, it provides the first evaluation of fMRI-fNIRS correspondence across multiple scales, including edge-, node-, graph-, and module-level analyses, enabling a nuanced characterization of where the modalities converge and diverge. A key methodological advantage is the fNIRS channel-to-spherical fMRI-ROI mapping, which allowed anatomically grounded regional comparisons. Further, near whole-head fNIRS coverage enabled a more complete assessment of RSN architecture, and modular analyses revealed meaningful subnetwork overlaps, marking the first systematic, quantitative demonstration of cross-modal similarity in community structure.

Despite using different cohorts, modality-specific preprocessing, and imperfect ROI-to-channel mapping, fNIRS and fMRI showed moderate agreement in global RSFC patterns, several graph metrics and modular organization. This convergence under suboptimal conditions suggests that true cross-modal correspondence may be even stronger, supporting the potential of fNIRS for characterizing large-scale functional networks.

Several limitations warrant consideration. First, different cohorts and independent preprocessing constrained precision and prevented within-subject comparison. Second, the fNIRS acquisition was shorter (5 vs. 15 min), potentially reducing RSFC reliability. Third, spatial uncertainty from channel-to-ROI mapping was substantial, as approximated positions using the fOLD toolbox led to the exclusion of 20 channels, limiting motor network coverage and cross-modal precision. Fourth, path-based graph metrics were restricted, with betweenness centrality excluded due to trivial values in fully connected networks, leaving only nodal path length for shortest-path analysis. Finally, larger, and more diverse samples are needed to establish robust population-level estimates.

These limitations primarily reflect dataset constraints rather than analytical weaknesses. Future work using larger cohorts, full-head coverage, individualized digitized channel positions, or ideally simultaneous fNIRS-fMRI recordings will improve anatomical precision, reproducibility and allow direct quantification of true cross-modal convergence, ultimately clarifying fNIRS translational utility for clinical RSFC tracking.

## 5 Conclusion

In summary, this study demonstrates that fNIRS captures key aspects of large-scale RSN organization observed with fMRI, particularly at the graph and module levels, while agreement is lower at finer scales such as edges and nodes. fNIRS reliably reproduced global connectivity patterns and major functional subnetworks, especially with HbO-based bivariate correlations. Nodal-level agreement was metric-dependent, with normalized nodal strength and assortativity being robust, whereas path-based and efficiency measures were more modality-specific. Partial correlations improved edge-level correspondence but reduced nodal and modular consistency. Overall, these results indicate that fNIRS is well suited for population-level characterization and longitudinal monitoring of RSN topology, while fine-grained or subject-specific connectivity estimates should be interpreted with caution. These findings position fNIRS as a promising, accessible and clinically relevant tool for non-invasive assessment of functional brain networks.

## 6 Glossary

(FC): functional connectivity
(fMRI): functional magnetic resonance imaging
(rs): resting-state
(RSFC): resting-state functional connectivity
(RSN): resting-state networks
(DMN): default mode network
(SMN): sensorimotor network
(CEN): central executive network
(AN): attentional network
(LN): language network
(VN): visual network
(SN): salience network
(fNIRS): near-infrared spectroscopy
(HbO): oxygenated hemoglobin
(HbR): deoxygenated hemoglobin
(HCP): Human Connectome Project
(BH FDR): Benjamini-Hochberg false discovery rate
(TOST): one-sided t-tests
(d-AN): dorsal-AN
(c-VN): central-VN
(p-VN): and peripheral-VN.

## 7 Data availability

The fMRI data analyzed in this study were obtained from the Human Connectome Project (HCP) and are publicly available via the HCP data repository (https://www.humanconnectome.org), subject to the HCP data use terms. The fNIRS data used in this study are part of a dataset currently under peer review for public release and are therefore not yet publicly available. These data can be obtained from the respective authors upon reasonable request. All analysis code, together with derived data and raw analysis outputs underlying the reported results, are available in an OSF repository (https://osf.io/w52v3/overview?view_only=a34c0de9d85c4b839639b97cf665f463). The repository includes a detailed README file that documents the analysis pipeline and provides step-by-step instructions to reproduce the results.

## 10 Author Contributions

**Foivos Kotsogiannis:** Conceptualization, Methodology, Software, Validation, Formal analysis, Investigation, Resources, Data Curation, Writing - Original Draft, Writing - Review & Editing, Visualization; **Michael Lührs**: Validation, Resources, Data Curation, Writing - Review & Editing; **Geert-Jan M. Rutten:** Validation, Writing - Review & Editing; **Andrew T. Reid:** Validation, Writing - Review & Editing; **Sabine Deprez**: Validation, Writing - Review & Editing; **Maarten Lambrecht:** Validation, Writing - Review & Editing; **Wouter De Baene:** Conceptualization, Validation, Writing - Original Draft, Writing - Review & Editing, Supervision; **Charlotte Sleurs:** Conceptualization, Methodology, Validation, Writing - Original Draft, Writing - Review & Editing, Supervision, Project administration, Funding acquisition

## 11. Acknowledgements

Data were provided [in part] by the Human Connectome Project, WU-Minn Consortium (Principal Investigators: David Van Essen and Kamil Ugurbil; 1U54MH091657) funded by the 16 NIH Institutes and Centers that support the NIH Blueprint for Neuroscience Research; and by the McDonnell Center for Systems Neuroscience at Washington University.

We also thank Bettina Sorger, Sophie Raible and João Pereira (Maastricht University) for their support and assistance in providing access to the fNIRS data.

## 13. Appendix A: Extended Methods

### 13.1 Edgewise similarity analysis: T-test & TOST

At the subject-level, an edgewise comparison was conducted. For each edge (ROI pair), two vectors were formed, containing Fisher’s *z* transformed correlation values across subjects for that edge in each modality:

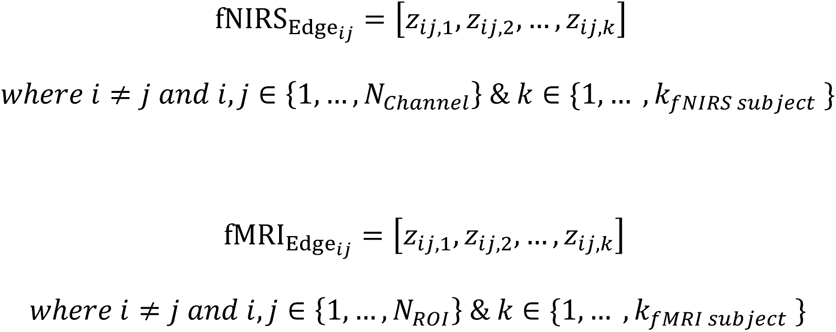

Because the fNIRS and fMRI groups are unpaired, after ensuring normal distribution of the edges, an independent two-sample’s *t*-test was conducted to assess the differences in connectivity strength between modalities. This procedure was repeated for all edges, producing matrices of *t*- and *p-*values. To account for multiple-comparisons, Benjamini-Hochberg false discovery rate (BH FDR) correction was performed on the upper triangle *p*-values.

While the *t*-test evaluates whether edges differ significantly between modalities, the non-significant results do not necessarily indicate similarity. Instead, they may reflect small differences or limited statistical power. Therefore, the two one-sided *t-*test (TOST) procedure (Schuirmann, 1981) for independent samples was conducted. This is an equivalence test in which a lower (−*ε*) and upper (+*ε*) equivalence bound is defined, and it tests two null hypotheses:

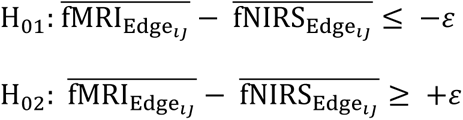

When both *t-*tests get rejected then the observed difference falls within the equivalence range 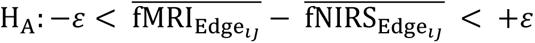 and we can conclude that the edge is considered statistically equivalent across modalities.

Because no prior work has defined bounds for cross-modal RSFC comparison, we explored a range of bounds (*ε* = 0.01 − 0.4, step = 0.001) to determine the interval in which the edges could be equivalent. The bounds were defined in terms of *r*-value differences and once the range of *r*-values was obtained it was then converted to Fisher *z*-scores to match the Fisher-*z* transformed edge data. TOST was then performed using the Fisher *z* edge values and the corresponding *z-*bounds. This procedure yielded a *p*-value TOST map, from which the upper triangular matrix (excluding the diagonal) was extracted and corrected for multiple comparisons using the BH FDR method.

### 13.2 Similarity Between Group-Averaged RSFC Matrices

At the group-level, similarity between fNIRS (HbO and HbR) and fMRI resting-state functional connectivity (RSFC) patterns was quantified by directly correlating their group-averaged connectivity matrices, combined with permutation testing to assess significance. Analyses were performed separately for matrices derived using Pearson’s correlation and partial correlation.

Specifically, for each modality and correlation approach, subject-level connectivity matrices were Fisher z-transformed and averaged across participants to obtain group-level matrices. To quantify cross-modal similarity, the upper triangular elements (excluding the diagonal) of the fMRI and fNIRS group-averaged matrices were vectorized and correlated using Pearson’s correlation coefficient. This yielded a single similarity metric reflecting the extent to which the relative pattern of stronger and weaker functional connections was preserved across modalities.

Because edges in connectivity matrices are not statistically independent, statistical significance was assessed using permutation testing. A null distribution was generated by randomly permuting the row and column labels of the fMRI connectivity matrix. For each permutation, the upper triangular elements of the permuted fMRI matrix were correlated with the fixed fNIRS matrix. This procedure was repeated 100,000 times, producing a null distribution of correlation coefficients expected under the hypothesis of no cross-modal RSFC similarity.

The permutation *p*-value was computed as the proportion of permuted correlations whose absolute value was equal to or greater than the observed correlation coefficient, using a two-sided test with a +1 correction to avoid zero p-values. This approach directly tests whether the observed similarity between fMRI and fNIRS RSFC patterns exceeds what would be expected by chance, while explicitly accounting for the non-independence of matrix entries.

### 13.3 Evaluation of Graph metrics

To further evaluate the similarity of functional connectivity between modalities, graph-theoretical metrics were computed on both subject and group-level correlation matrices, treated as undirected weighted graphs. All analyses were performed separately for bivariate and partial correlation matrices.

To ensure valid comparison of graph metrics across modalities, the correlation matrices were normalized to have comparable amplitude properties. For both fNIRS (HbO and HbR) and fMRI, each correlation matrix was first converted to Fisher *z* values (for both individual- and group-level matrices). Then, the absolute maximum Fisher *z* value (*abs*_*max*_ = max(|*z*|)) across all matrices was identified (separately for bivariate and partial matrices). Each matrix was divided by this global maximum to ensure consistent scaling while preserving zero correlations.

Graph-theoretical metrics were estimated using BCTpy (Rubinov et al., 2009; Rubinov and Sporns, 2010). The metrics included nodal strength, assortativity, nodal efficiency, betweenness centrality, and nodal path length. Given the use of signed connectivity matrices, two complementary measures of nodal strength were computed. Net nodal strength, defined as the sum of all positive and negative edge weights, and normalized nodal strength, defined as a weighted average of positive and negative edges which is a more representative estimate for signed networks (Fornito et al., 2016b; Rubinov and Sporns, 2011) which aligns more closely with neurophysiology (Rubinov and Sporns, 2011). To estimate betweenness centrality and nodal path length, correlation matrices were converted into distance matrices (Fornito et al., 2016b; Rubinov et al., 2009; Rubinov and Sporns, 2010). Specifically, *z*-transformed correlation matrices were first converted to absolute values to retain the physiological relevance of both positive (excitatory) and negative (inhibitory) connections (Fornito et al., 2016b). These absolute |*z*| were then converted to distances *d*_*i,j*_ = 1 − |*z*_*ij*_|. This ensured that stronger functional connectivity corresponded to shorter distances. For nodal efficiency, edge weights needed to be in the range of 0 to 1; therefore, the absolute values from both individual and group matrices were used.

#### 13.3.1 Subject-level analyses

On the subject-level, nodal differences in graph metrics were assessed using independent samples *t-*tests. For each node, the graph metric of interest (e.g. net nodal strength) was computed separately for every subject in each modality. This produced for each node *i, where i* ∈ {*ROI*1, *ROI*2, …, *ROI*82}, a modality specific vector containing the metric values across subjects (e.g. one vector for fMRI subjects and one vector for fNIRS subjects). These vectors were then compared between modalities using independent samples *t*-test to determine whether the nodal graph properties differed significantly. Multiple comparisons across nodes were corrected with the BH FDR method.

#### 13.3.2 Group-level analyses

To quantify the similarity of the graph metrics derived from group-averaged matrices across modalities, the spatial distribution of each nodal metric was correlated between fMRI and fNIRS. For each metric, a vector containing the nodal values across all ROIs was formed and correlated with the corresponding vector from the other modality. This procedure quantifies the similarity in the topological organization of the nodal graph properties between fMRI and fNIRS. The correlation statistic (i.e. Pearson’s *r*, Spearman’s *rho*) was selected based on Shapiro-Wilk tests evaluating whether the nodal distributions met the assumptions for parametric testing.

### 13.4 Modularity Analysis & Network Overlap

To further evaluate the graph properties of the group matrices across modalities, modularity was estimated using the Louvain algorithm. Due to the stochastic nature of the algorithm, to ensure stable community assignments multiple runs were performed followed by consensus clustering.

Specifically, a range of resolution parameters *γ* from 0.3 to 2.0 were used (lower *γ* yields larger communities, while higher smaller communities). For each *γ*, 1,000 iterations of the algorithm were performed. Then the agreement matrix was computed for every node pair, and finally a clustering algorithm was applied to extract the most robust, representative community structure (Bassett et al., 2013; Sporns and Betzel, 2016). This produces a single, reliable community assignment that best reflects the repeated stochastic community assignments.

Finally, community assignments producing five to seven modules were considered to identify the most biologically plausible organization of ROIs. The gamma value producing seven communities was chosen after visually inspecting the communities overlaid on RSN templates (Yeo et al., 2011) and their MNI coordinates were confirmed to correspond with functionally relevant rs-fMRI networks using NeuroSynth (Yarkoni et al., 2011).

For each modality, the final community assignments (with seven stable unique communities) were first converted into sets, where each community label represented the collection of ROIs assigned to the module. For each fMRI community 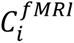, its similarity to every fNIRS community 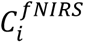 was computed using Jaccard similarity:

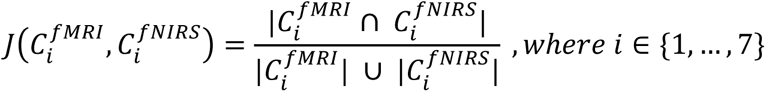

which quantifies the proportion of shared ROIs for each community label between modalities. For each fMRI community, the fNIRS community that yielded the maximum Jaccard coefficient was identified as its optimal match. This was performed separately for HbO and HbR community structures and for both bivariate and partial correlation approaches to estimate FC. Finally, an overall correspondence index was computed by averaging the maximum Jaccard coefficients across all seven communities, providing a single summary value of total fMRI-fNIRS community overlap.

To assess whether the observed correspondence between fMRI and fNIRS communities was statistically significant, a non-parametric permutation test was performed. After identifying the best overlapping fNIRS community for each fMRI community, a null distribution was generated by randomly shuffling the fNIRS ROIs and reassigning them to communities while keeping the original fMRI community structure fixed. For each permutation, the Jaccard index was estimated between the permuted fNIRS communities and the fixed fMRI communities, and the maximum Jaccard value for each fMRI community was extracted. This procedure was repeated 100,000 times, yielding a null distribution of maximum Jaccard values expected by chance. To obtain a *p*-value for each fMRI community, its observed maximum Jaccard value was compared against its corresponding null-distribution.

### 13.5 Module RSN labeling

For the resulting seven-stable fMRI modules, network identity was assigned using a data-driven approach based on the 17-network canonical RSNs from Yeo et al. (2011). ROI coordinates in MNI space were overlaid onto the canonical RSN maps to quantify spatial correspondence. Each module consisted of nodes defined by 5mm radius spheres, and for each node, the number of voxels overlapping with each of the 17 RSNs was computed. Module RSN labeling was then determined by summing voxel overlaps across all nodes within a module and the network label was assigned based on the highest voxel count.

In some cases, multiple RSN labels were assigned to a single module, reflecting a mixed network composition. This was reported when substantial voxel overlap was observed with more than one RSN. For example, as shown in Table 3, bivariate fMRI module 6 overlaps predominantly with both c-VN (44% of voxels) and with p-VN (33% of voxels). Detailed voxel-wise overlap proportions of modules are provided in Table B.1.

**Table B.1.**
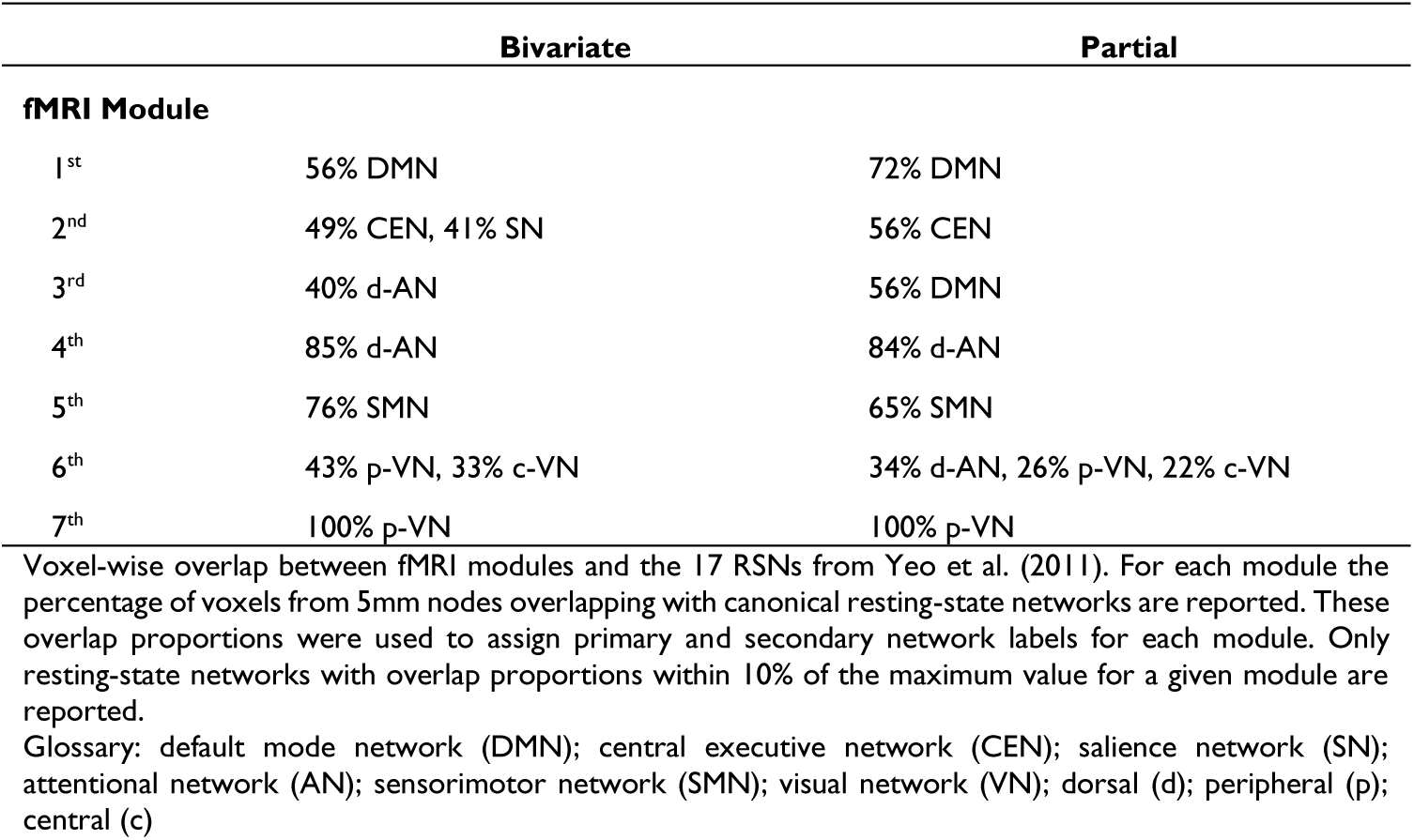
Fraction of module voxels assigned to each canonical RSN.

## 14 Appendix B: Additional Figures

**Figure B.1.**
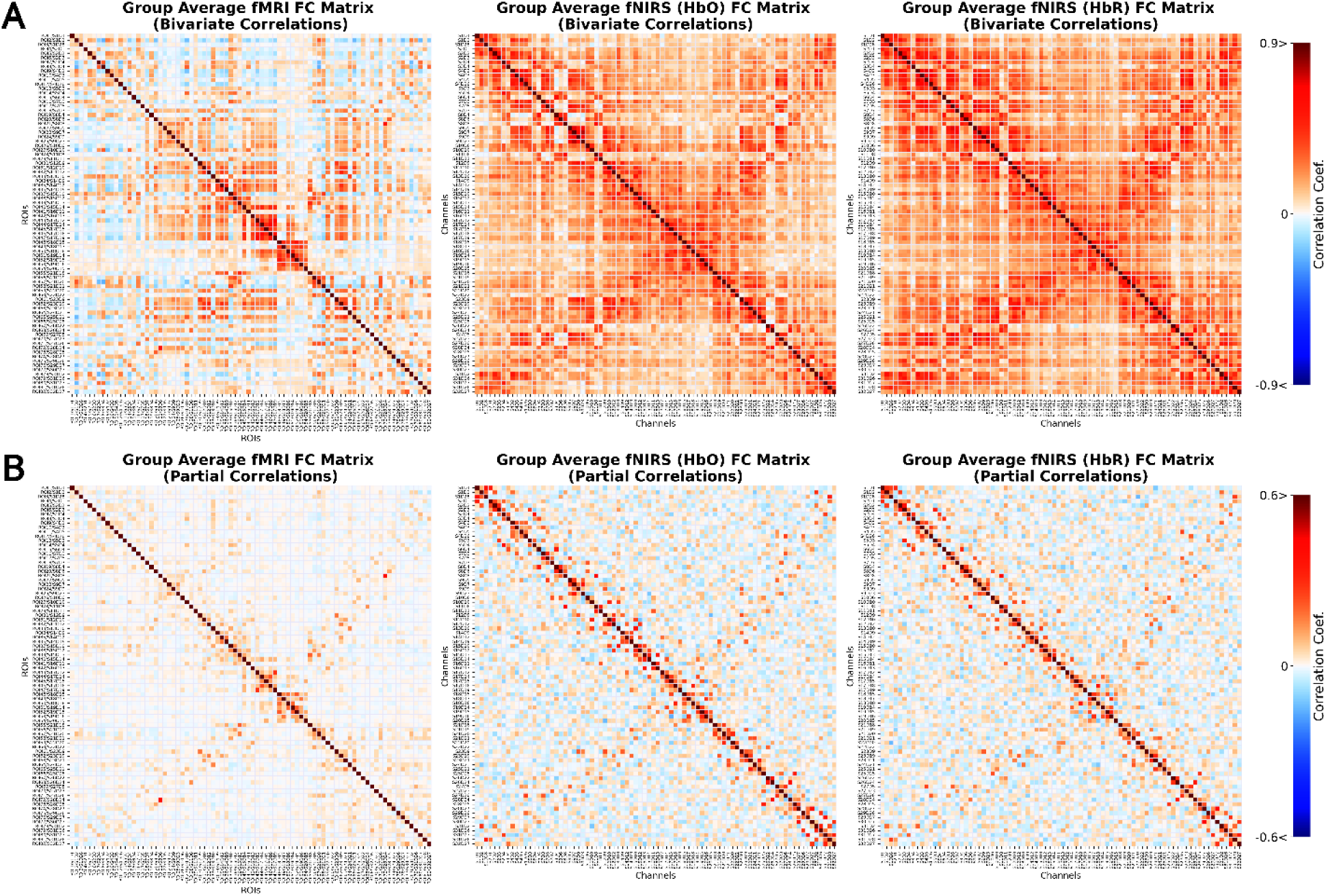
Group-level RSFC matrices. Panel (A) represents FC matrices derived from bivariate and panel (B) from partial correlations. Matrices are ordered according to the original channel naming scheme. The ordering between fMRI ROIs and fNIRS channels is aligned based on the ROI-channel mapping. Results are shown for fMRI, fNIRS-HbO and fNIRS-HbR.

**Figure B.2.**
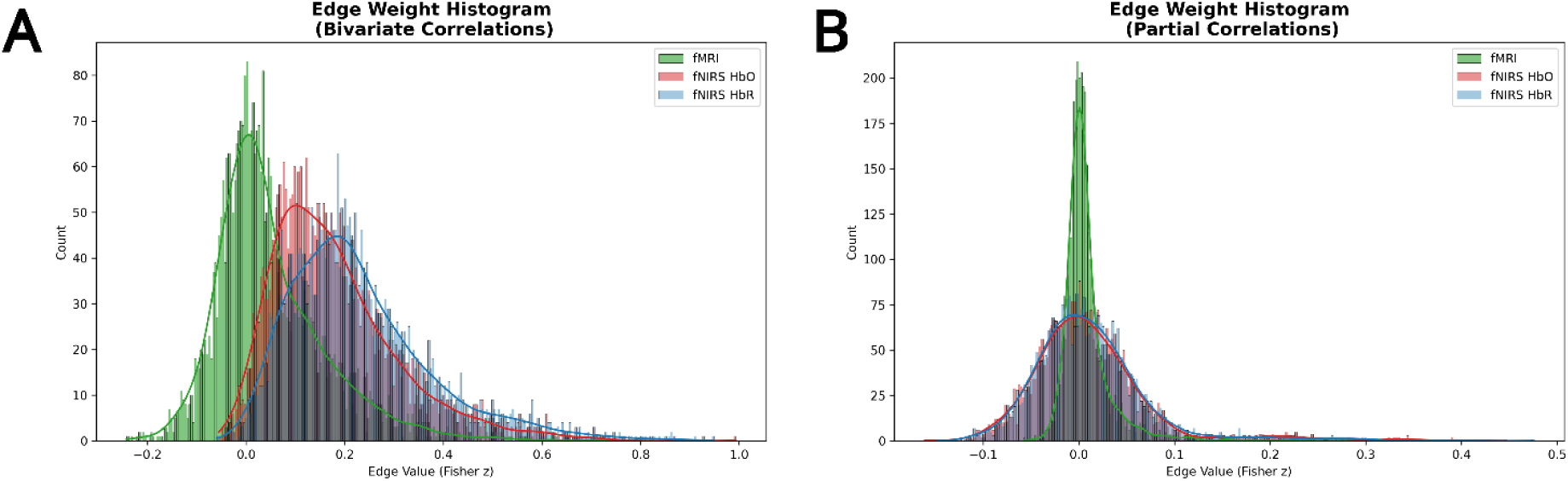
Distribution of edge values for fMRI, fNIRS-HbO and fNIRS-HbR excluding diagonal and lower-triangular elements. Panel (A) shows the distribution of average bivariate correlation matrices, while Panel (B) illustrates the distribution of average partial correlation matrices. Histograms represent edge counts, and the overlaid lines the estimated distribution calculated from kernel density estimation.

**Figure B.3.**
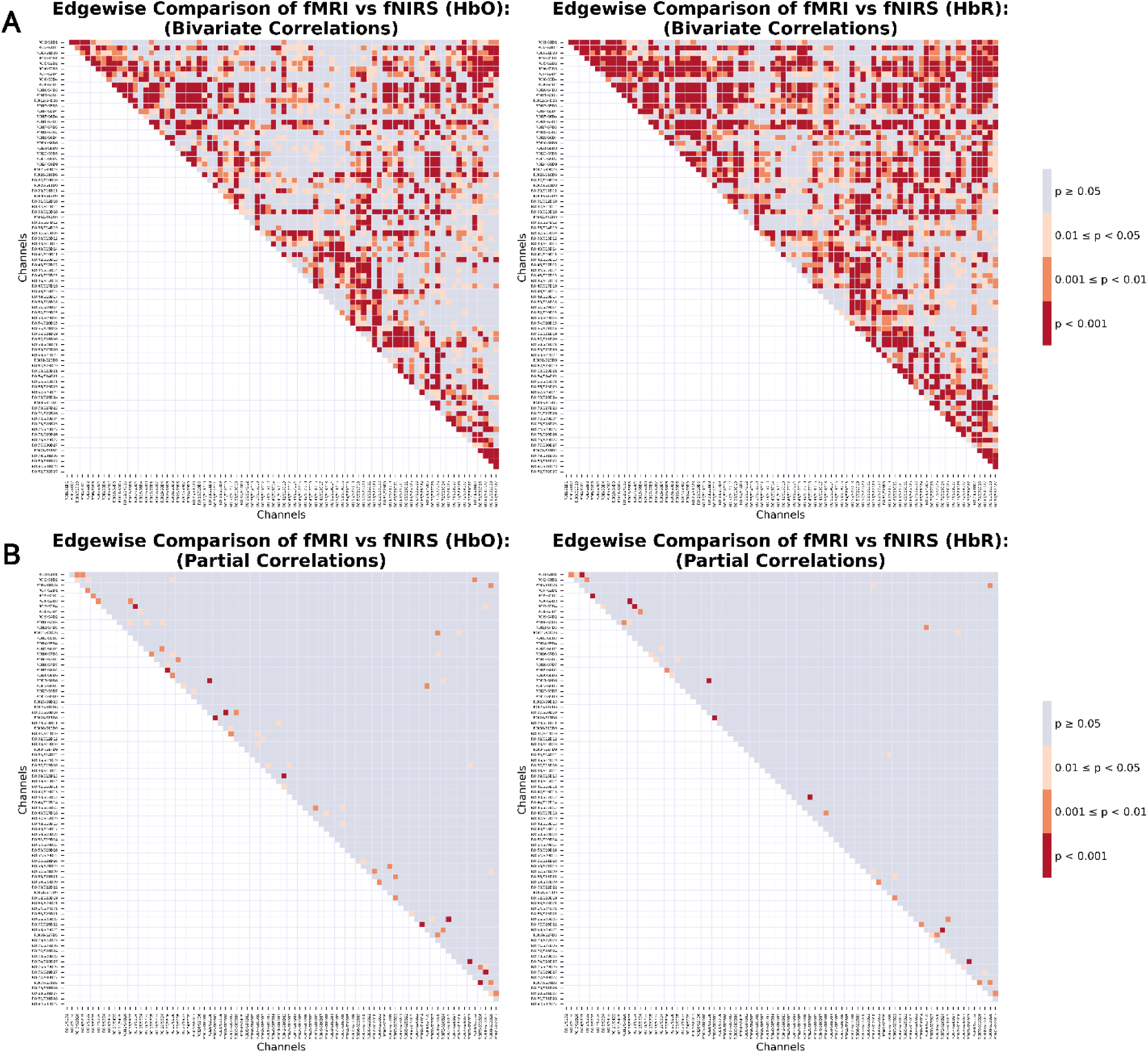
Edgewise differences. Independent sample’s *t-*test *p*-map (BH FDR corrected) on edges across subjects between modalities. Panel (A) illustrates the *t*-test results from bivariate correlation RSFC, while panel (B) from partial correlation RSFC.

**Figure B.4.**
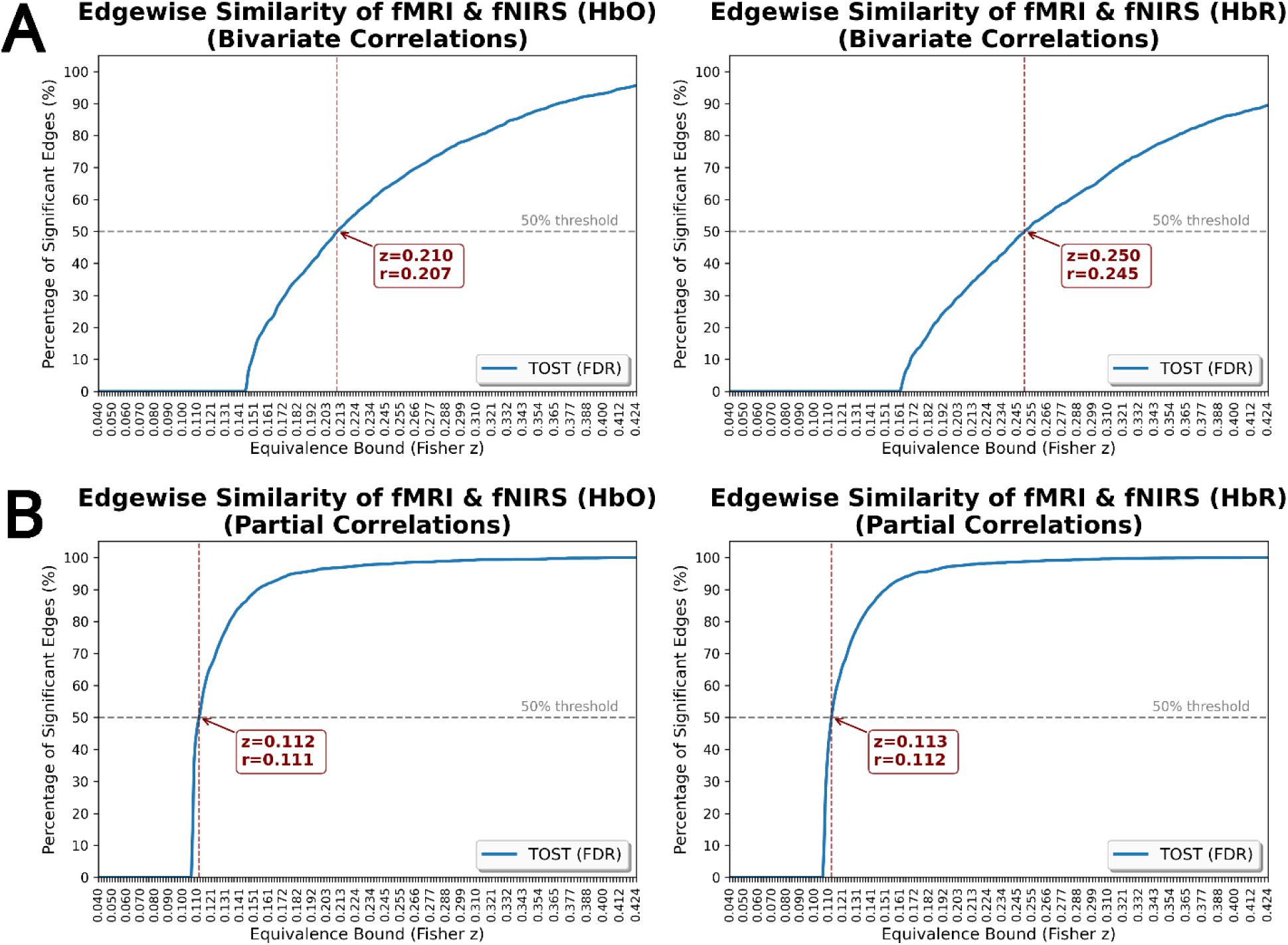
Edgewise equivalence bounds. Edgewise TOST results from (A) bivariate correlation matrices and (B) partial correlation matrices. The range of equivalence bounds is defined as the correlation *r* range of 0.01 to 0.4 (step = 0.001) converted to Fisher’s *z* to match the edge data. For each TOST *p-*map BH FDR correction was applied on the upper triangular portion of the matrix.

