## Supplementary Materials for "Light on Broken Networks: Resting-State fNIRS as a Tool for Connectivity Mapping"

### Supplementary Materials & Figures

#### Extended Method Section

##### Edgewise similarity analysis: T-test & TOST

At the subject-level, an edgewise comparison was conducted. For each edge (ROI pair), two vectors were formed, containing Fisher's *z* transformed correlation values across subjects for that edge in each modality:

$$\text{fNIRS}_{\text{Edge}_{ij}} = \left[z_{ij,1}, z_{ij,2}, \dots, z_{ij,k}\right]$$

$$where\ i \neq j\ and\ i, j \in \{1, \dots, N_{Channel}\}\ \&\ k \in \{1, \dots, k_{fNIRS\ subject}\}$$

$$\text{fMRI}_{\text{Edge}_{ij}} = \left[z_{ij,1}, z_{ij,2}, \dots, z_{ij,k}\right]$$

$$where\ i \neq j\ and\ i, j \in \{1, \dots, N_{ROI}\}\ \&\ k \in \{1, \dots, k_{fMRI\ subject}\}$$

Because the fNIRS and fMRI groups are unpaired, after ensuring normal distribution of the edges, an independent two-sample's *t*-test was conducted to assess the differences in connectivity strength between modalities. This procedure was repeated for all edges, producing matrices of *t*- and *p*-values. To account for multiple-comparisons, Benjamini-Hochberg false discovery rate (BH FDR) correction was performed on the upper triangle *p*-values.

While the *t*-test evaluates whether edges differ significantly between modalities, the non-significant results do not necessarily indicate similarity. Instead, they may reflect small differences or limited statistical power. Therefore, the two one-sided *t*-test (TOST) procedure (Schuirmann, 1981) for independent samples was conducted. This is an equivalence test in which a lower $(-\varepsilon)$ and upper $(+\varepsilon)$ equivalence bound is defined, and it tests two null hypotheses:

$$\text{H}_{01}\text{: } \overline{\text{fMRI}_{\text{Edge}_{ij}}} - \overline{\text{fNIRS}_{\text{Edge}_{ij}}} \leq -\varepsilon$$

$$\text{H}_{02}\text{: } \overline{\text{fMRI}_{\text{Edge}_{ij}}} - \overline{\text{fNIRS}_{\text{Edge}_{ij}}} \geq +\varepsilon$$

When both $t$-tests get rejected then the observed difference falls within the equivalence range $\mathrm{H_A}: -\varepsilon < \overline{\mathrm{fMRI}_{\mathrm{Edge}_{ij}}} - \overline{\mathrm{fNIRS}_{\mathrm{Edge}_{ij}}} < +\varepsilon$ and we can conclude that the edge is considered statistically equivalent across modalities.

For each modality, the final community assignments (with seven stable unique communities) were first converted into sets, where each community label represented the collection of ROIs assigned to the module. For each fMRI community $C_i^{fMRI}$, its similarity to every fNIRS community $C_i^{fNIRS}$ was computed using Jaccard similarity:

**Table 1. Fraction of module voxels assigned to each canonical RSN**

| | Bivariate | Partial |
|---|---|---|
| **fMRI Module** | | |
| 1st | 56% DMN | 72% DMN |
| 2nd | 49% CEN, 41% SN | 56% CEN |
| 3rd | 40% d-AN | 56% DMN |
| 4th | 85% d-AN | 84% d-AN |
| 5th | 76% SMN | 65% SMN |
| 6th | 43% p-VN, 33% c-VN | 34% d-AN, 26% p-VN, 22% c-VN |
| 7th | 100% p-VN | 100% p-VN |

Voxel-wise overlap between fMRI modules and the 17 RSNs from Yeo et al. (2011)[22]. For each module the percentage of voxels from 5mm nodes overlapping with canonical resting-state networks are reported. These overlap proportions were used to assign primary and secondary network labels for each module. Only resting-state networks with overlap proportions within 10% of the maximum value for a given module are reported.

**Abbreviations**: default mode network (DMN); central executive network (CEN); salience network (SN); attentional network (AN); sensorimotor network (SMN); visual network (VN); dorsal (d); peripheral (p); central (c)

### Supplementary Figures

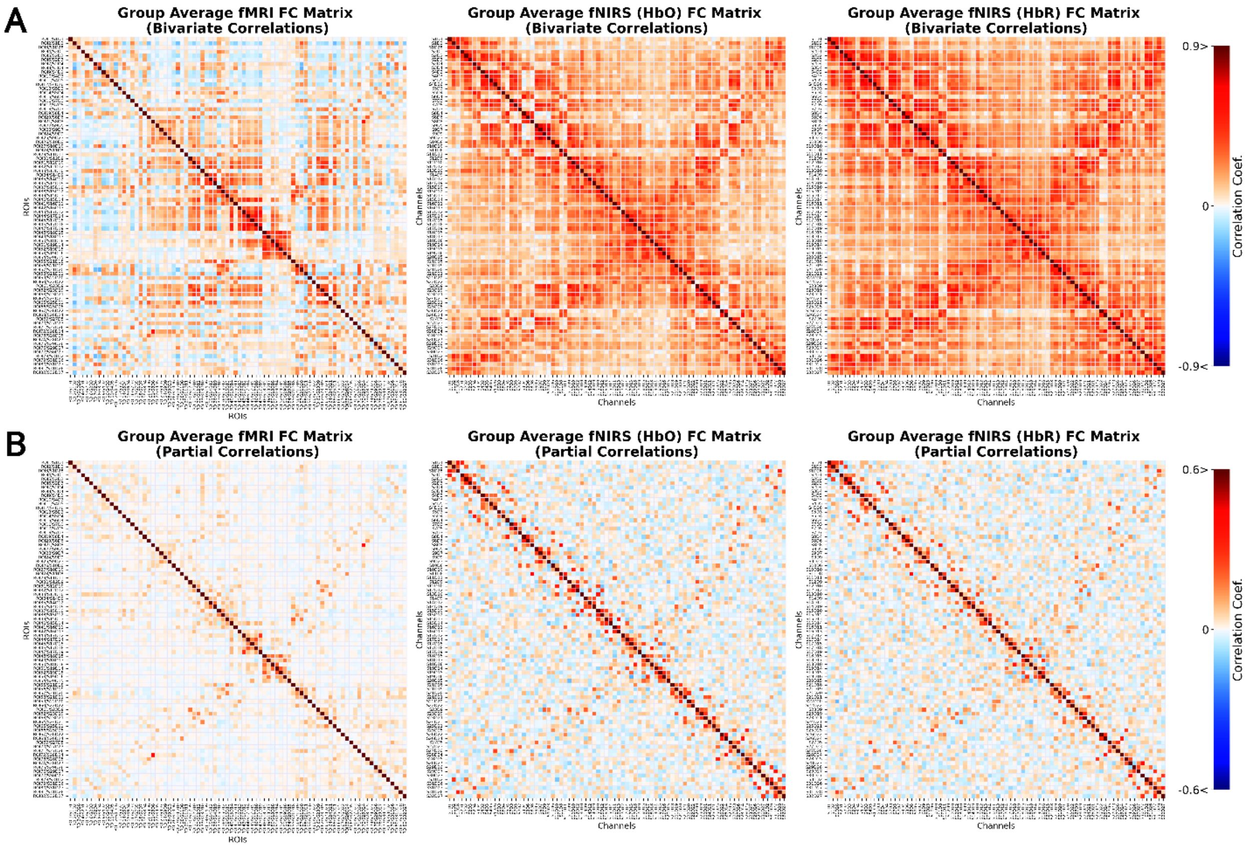


**Figure 1 Group-level RSFC matrices**. Panel (A) represents FC matrices derived from bivariate and panel (B) from partial correlations. Matrices are ordered according to the original channel naming scheme. The ordering between fMRI ROIs and fNIRS channels is aligned based on the ROI-channel mapping. Results are shown for fMRI, fNIRS-HbO and fNIRS-HbR.

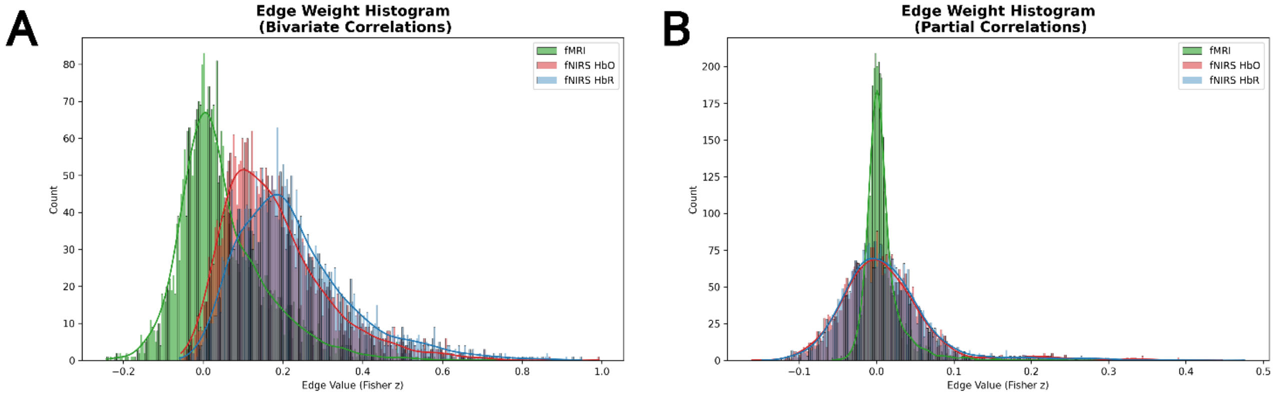


**Figure 2 Distribution of edge values for fMRI, fNIRS-HbO and fNIRS-HbR excluding diagonal and lower-triangular elements**. Panel (A) shows the distribution of average bivariate correlation matrices, while Panel (B) illustrates the distribution of average partial correlation matrices. Histograms represent edge counts, and the overlaid lines the estimated distribution calculated from kernel density estimation.

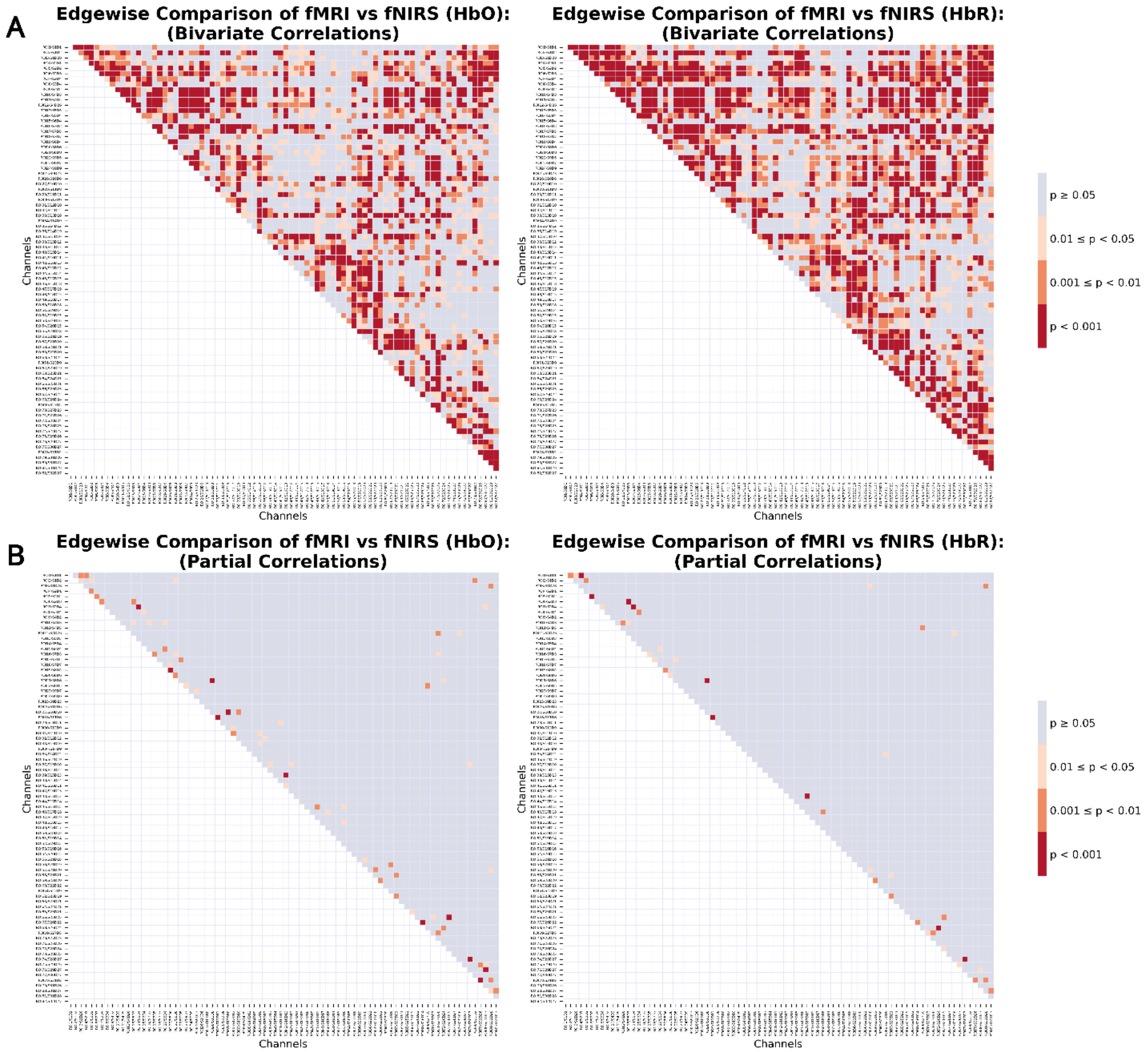


**Figure 3 Edgewise differences.** Independent sample's *t*-test *p*-map (BH FDR corrected) on edges across subjects between modalities. Panel (A) illustrates the *t*-test results from bivariate correlation RSFC, while panel (B) from partial correlation RSFC.

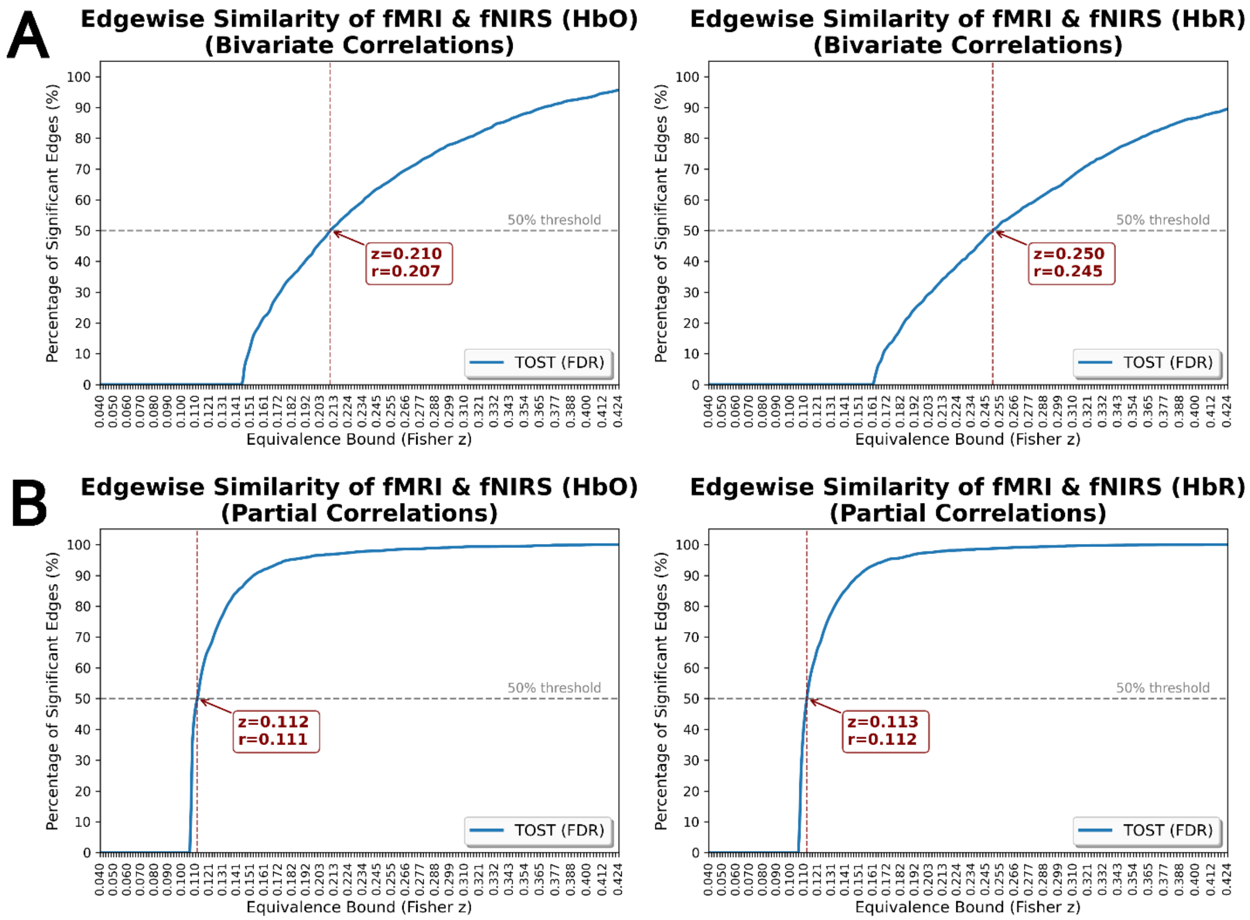


**Figure 4 Edgewise equivalence bounds.** Edgewise TOST results from (A) bivariate correlation matrices and (B) partial correlation matrices. The range of equivalence bounds is defined as the correlation *r* range of 0.01 to 0.4 (step = 0.001) converted to Fisher's *z* to match the edge data. For each TOST *p*-map BH FDR correction was applied on the upper triangular portion of the matrix.
